# Interbacterial competition and anti-predatory behavior of environmental *Vibrio cholerae* strains

**DOI:** 10.1101/2020.07.24.220178

**Authors:** Natália C. Drebes Dörr, Melanie Blokesch

## Abstract

*Vibrio cholerae* isolates responsible for cholera pandemics represent only a small portion of the diverse strains belonging to this species. Indeed, most *V. cholerae* are encountered in aquatic environments. To better understand the emergence of pandemic lineages, it is crucial to discern what differentiates pandemic strains from their environmental relatives. Here, we studied the interaction of environmental *V. cholerae* with eukaryotic predators or competing bacteria and tested the contributions of the hemolysin and the type VI secretion system (T6SS) to those interactions. Both of these molecular weapons are constitutively active in environmental isolates but subject to tight regulation in the pandemic clade. We showed that several environmental isolates resist amoebal grazing and that this anti-grazing defense relies on the strains’ T6SS and its actin-cross-linking domain (ACD)-containing tip protein. Strains lacking the ACD were unable to defend themselves against grazing amoebae but maintained high levels of T6SS-dependent interbacterial killing. We explored the latter phenotype through whole-genome sequencing of fourteen isolates, which unveiled a wide array of novel T6SS effector and (orphan) immunity proteins. By combining these *in silico* predictions with experimental validations, we showed that highly similar but nonidentical immunity proteins were insufficient to provide cross-immunity among those wild strains.

**Originality-Significance Statement:** This work contributes to the understanding of phenotypic consequences that differentiate diverse *Vibrio cholerae* strains. We focused on the type VI secretion system (T6SS) and the pore forming toxin hemolysin, which are tightly regulated in pandemic strains but remain constitutively active in non-pandemic isolates. We unveiled diverse arrays of T6SS effector/immunity modules in a set of environmental strains by long-read whole genome sequencing and *de novo* assembly. These modules determine whether the strains are able to evade amoebal predation and dictate their level of compatibility or competitiveness with one another.

## Introduction

Diarrheal diseases can be caused by a variety of microorganisms, including the causative agent of cholera, *Vibrio cholerae*, which infects up to 4 million people every year (Ali *et al*., 2015). Cholera often spreads from its endemic area around the Ganges delta and has reached almost the entire world in the reported seven pandemics that have been witnessed since 1817. In addition to these pandemics, important localized outbreaks have occurred over the years, especially following natural disasters (Faruque *et al*., 1998; World Health Organization (WHO), 2006; Clemens *et al*., 2017).

*V. cholerae* strains can be classified into serogroups based on more than 200 different O-antigens. The O1 serogroup is considered the primary cause of previous (e.g., classical serotype) and ongoing (e.g., El Tor serotype) cholera pandemics (Faruque *et al*., 1998; Cottingham *et al*., 2003). Isolates belonging to the O139 serogroup are genetically related to the 7^th^ pandemic O1 El Tor strains (Johnson *et al*., 1994) but are rarely associated with disease outbreaks currently (Faruque *et al*., 1998; Clemens *et al*., 2017). Even though this species is best known due to the life-threatening disease it causes, the vast majority of *V. cholerae* are common members of aquatic habitats. These mostly non-O1/non-O139 serogroup strains are thought to frequently associate with zooplankton and shellfish, and *V. cholerae* uses their chitinous exoskeletons as a source of carbon and nitrogen (Faruque *et al*., 1998; Cottingham *et al*., 2003; Kirn *et al*., 2005). Environmental isolates are typically considered harmless to humans despite reported associations with mild to severe forms of diarrhea (Islam *et al*., 1992; Singh *et al*., 2001; Dziejman *et al*., 2002, 2005; Onifade *et al*., 2011; Deshayes *et al*., 2015; Hasan *et al*., 2015).

A major question, not only in the cholera field but also for infectious diseases in general, is how pathogenic isolates evolve from their non-pathogenic environmental progenitors. In the case of *V. cholerae*, pandemic patient isolates are remarkably clonal, in sharp contrast to the high genomic variability encountered in environmental isolates (Faruque *et al*., 1998; Chun *et al*., 2009; Mutreja *et al*., 2011; Harris *et al*., 2012; Domman *et al*., 2017; Weill *et al*., 2017, 2019). Two major genetic features common to all pandemic strains are the CTXΦ prophage and the *Vibrio* pathogenicity island (VPI-1 or TCP island). These genetic elements harbor genes encoding the main virulence factors, e.g., cholera toxin (CTX) and toxin-coregulated pilus (TCP) (Taylor *et al*., 1987; Waldor and Mekalanos, 1996). The possession of these genomic regions, however, is not unique nor a ‘deterministic factor’ of pandemic strains. In fact, *V. cholerae* samples collected in several regions of the world have exposed environmental CTX- and/or TCP-positive strains (Rivera *et al*., 2001; Faruque *et al*., 2003, 2004; Gennari *et al*., 2012; Bernardy *et al*., 2016; Shapiro *et al*., 2016). Additionally, *V. cholerae* strains that have caused localized cholera outbreaks without reaching pandemic levels have been reported, such as the O37 serogroup strains V52 and ATCC25872 (Aldova *et al*., 1968; Boyd and Waldor, 2002; Cottingham *et al*., 2003; Chun *et al*., 2009). Therefore, the terms “toxigenic” (ability to cause cholera) and “pandemic” (here, current 7^th^ pandemic-causing O1 El Tor strains) are not synonymous. What exactly determines whether strains become pandemic or not is still not fully understood and probably involves diverse aspects ranging from genetic content and strain-specific phenotypes, along with the location of the initial outbreaks and ultimately towards social and sanitary factors, to name a few. Pertinent to the bacteria-related aspects, previous studies hypothesized the existence of virulence adaptive polymorphisms (VAP) circulating in environmental strains. These VAPs were suggested to be a prerequisite for pandemic transition before the horizontal acquisition of CTXΦ and VPI-1 (Shapiro *et al*., 2016).

To better understand their emergence, the differences between pandemic strains and their environmental relatives need to be deciphered. Such differences most likely include phenotypic alterations that are not easily predictable by genomics. In the context of phenotypic variation, two minor virulence factors, namely, the type VI secretion system (T6SS) and the pore-forming toxin hemolysin, are of special interest as they are differentially produced in pandemic compared with non-pandemic toxigenic or environmental strains. The T6SS is a molecular killing device that resembles an inverted contractile bacteriophage tail and it is present in approximately 25% of all Gram-negative bacteria (Ho *et al*., 2014; Cianfanelli *et al*., 2016; Galán and Waksman, 2018; Taylor *et al*., 2018). It is composed of a membrane-spanning portion, a tube structure made of stacks of Hcp hexamer rings, and a tube-surrounding sheath. The sheath is composed of the two proteins VipA and VipB which, upon contraction, propels the inner tube out of the cell together with its effector-decorated tip proteins (VgrG and PAAR) (Zoued *et al*., 2014; Cherrak *et al*., 2019; Flaugnatti *et al*., 2020). Secreted Hcp therefore serves as an indicator of T6SS activity (Pukatzki et al 2006; Basler *et al*., 2012; Bröms *et al*., 2013; Kube *et al*., 2014). The secreted effector proteins mostly target conserved cellular components, such as membranes, bacterial peptidoglycan, nucleic acids, or the eukaryotic cytoskeleton (Hood *et al*., 2010; Russell *et al*., 2014). T6SS effectors can be either secreted as (i) cargos that interact with T6SS structural proteins, such as Hcp or the tip protein VgrG; or as (ii) C-terminal extensions of VgrG, Hcp, or PAAR (so-called “evolved” proteins; Pukatzki *et al*., 2006, 2009; Hachani *et al*., 2016). Notably, T6SS-producing bacteria protect themselves against their toxic effector repertoire by the production of effector-cognate immunity proteins. These effector/immunity (E/I) pairs are usually encoded adjacent to each other (Russell *et al*., 2011; Brooks *et al*., 2013; Dong *et al*., 2013; Unterweger *et al*., 2014).

In current pandemic strains, the T6SS is encoded by four gene clusters: the large cluster and three auxiliary clusters. The large cluster primarily codes for structural proteins (recently reviewed by Crisan and Hammer, 2020), including the evolved tip protein VgrG3, which possesses a C-terminal lysozyme-like domain for peptidoglycan degradation (Zheng *et al*., 2011; Brooks *et al*., 2013). Auxiliary clusters 1 and 2 are both composed of genes encoding Hcp and VgrG (evolved in cluster 1 and structural in cluster 2), an adaptor protein (Tap1 or VasW; Liang *et al*., 2015; Unterweger *et al*., 2015) and an E/I module. The auxiliary cluster 1 effector TseL is a bifunctional lipase with anti-bacterial and anti-eukaryotic activity (Zheng *et al*., 2011; Dong *et al*., 2013; Russell *et al*., 2013). The auxiliary cluster 2 effector VasX acts as a pore-forming toxin due to its bacterial/eukaryotic colicin-like membrane-disrupting activity (Miyata *et al*., 2011, 2013; Russell *et al*., 2014). Finally, the T6SS auxiliary cluster 3 is composed of genes encoding a second copy of PAAR (a tip-sharpening protein that extends from VgrG; the first gene copy is at the start of the large cluster; Shneider *et al*., 2013) and a single E/I pair. When discovered, the aux 3 effector TseH was predicted to contain a hydrolase domain (Altindis *et al*., 2015), while the recently reported crystal structure supported its role as a papain-like NlpC/P60 peptidase (Hersch *et al*., 2020) with structural similarity to the T6SS effector Tse1 of *Pseudomonas aeruginosa* that contains bacteriolytic peptidoglycan amidase activity (Chou *et al*., 2012).

Another toxic protein that shows differential activity in pandemic compared with non-pandemic *V. cholerae* strains is the hemolysin protein (HlyA), which is likewise widespread among *Vibrio* species (Zhang and Austin, 2005). HlyA is considered a minor virulence factor as it contributes to toxicity in the context of intestinal infections (Ichinose *et al*., 1987; Olivier *et al*., 2007). We previously showed that the proper timing of HlyA activity is a prerequisite for pandemic *V. cholerae* to establish a replication niche within the aquatic amoeba *Acanthamoeba castellanii*, while constitutive activity kills this host prematurely (Van der Henst *et al*., 2018).

In this study, we deciphered phenotypic and genotypic differences between O1 El Tor pandemic strains and 15 environmental *V. cholerae* isolates. With respect to phenotypes, we focused primarily on T6SS- and hemolysin-specific outcomes exerted on competing bacteria or amoebal predators. We show that clade-specific anti-amoebal toxicity is dependent on the actin-cross-linking domain of a subset of T6SS effector proteins and that HlyA does not harm these predators under the tested conditions. Anti-bacterial activity, on the other hand, is widespread among the environmental isolates and is based on their constitutive T6SS production, which is a major phenotypic difference compared with the exquisitely regulated T6SS of pandemic *V. cholerae*. We also performed long-read PacBio whole-genome sequencing of the environmental isolates, which allowed us to perform basic comparative genomic analyses. Through this approach, we identified T6SS E/I modules as well as orphan immunity loci. Finally, we experimentally tested how the environmental isolates compete with one another and how this interbacterial competition correlates with their T6SS E/I repertoire.

## Results and discussion

### Genome sequencing of environmental *V. cholerae* strains

Apart from the major virulence factors, previous observations suggested important phenotypic differences between pandemic and environmental *V. cholerae* strains. We therefore decided to study fifteen environmental isolates (Table S1) from diverse habitats along the central California coast, a region that is free of endemic or epidemic cholera. These strains were initially isolated in 2004 by Keymer and colleagues followed by basic characterization and comparative genome hybridization (CGH) analyses (Keymer *et al*., 2007; Miller *et al*., 2007). The latter approach was based on amplicon microarrays (representing 3,357 of 3,891 annotated open reading frames), which were designed using the first published *V. cholerae* genome sequence as a template, namely, pandemic O1 El Tor strain N16961 (Heidelberg *et al*., 2000). Based on the presence or absence of the microarray-templated genes, the strains were classified into four clades (A-D) (Keymer *et al*., 2007; Miller *et al*., 2007), a classification we maintained throughout the current report. To better understand the accessory genome, including the T6SS E/I modules and to identify those genes that are novel when compared with strain N16961, we first whole-genome sequenced these strains using a long-read PacBio approach followed by the *de novo* assembly of their genomes. As a representative strain of the 7^th^ cholera pandemic, we used strain O1 El Tor A1552 throughout this study (Yildiz and Schoolnik, 1998). This strain is connected to a cholera outbreak in Peru in the 1990s. We recently reported its genome sequence, including more than 1,000 manual gene annotations, according to previous experimental validations (Matthey *et al*., 2018).

Table S2 shows the sequencing details and the features of the closed genomes of the 15 environmental isolates. All genomes showed the dual chromosome architecture that is common for *Vibrio* species (Okada *et al*., 2005), a similar overall size of the two chromosomes, and average GC percentages within the same range as the one observed for the pandemic *V. cholerae* strains N16961 and A1552 (Heidelberg 2000; Matthey *et al*., 2018). The assembly pipeline also predicted megaplasmids of ~300kbp and 80kbp for four strains belonging to clades C and D, respectively. It should be noted that due to the size selection of the prepared sequencing libraries, putative smaller plasmids remained unidentified. Interestingly, we observed that the genomes of strains W6G and W7G were almost identical. This reflects the previous report by Keymer *et al*. In their original sampling study, the authors claimed that, based on CGH, 30 unique genotypes were identified within their collection of 41 environmental strains, while several genotypes were sampled multiple times from distinct sampling events. Indeed, upon direct comparison of the W6G and W7G genomes, we observed identities of 99.98%, 99.997% and 99.998% for chromosome 1, chromosome 2, and the megaplasmid, respectively. Both strains possessed the same genes in all three replicons, and the few observed differences were primarily single nucleotides indels (which might, in part, reflect sequencing artifacts). These data suggest almost clonality between these two environmental samples (W6G and W7G). In addition, our whole-genome sequencing data showed that strains E7G and SA7G of clade D also had high levels of identity, which again confirmed the previous CGH data (Keymer *et al*., 2007). The majority of other strains differed more significantly and therefore allowed us to test links between specific phenotypes and the corresponding genotypes.

### Only a subset of environmental isolates block amoebal predation

We initially wondered how these environmental isolates would behave when confronted with predatory grazers, such as bacterivorous amoebae. We therefore tested the representative 7^th^ pandemic strain A1552 and the above-described collection of environmental strains for their ability to defend themselves against the grazing soil amoeba *Dictyostelium discoideum*. As shown in Figure 1A, a clade-specific behavior was observable in which strains from clade A and B appeared as non-toxic to *D. discoideum* and were efficiently grazed on while clade C and D strains completely resisted amoebal grazing. We concluded that anti-amoebal defenses significantly vary among the different environmental *V. cholerae* isolates.

**Figure 1:**
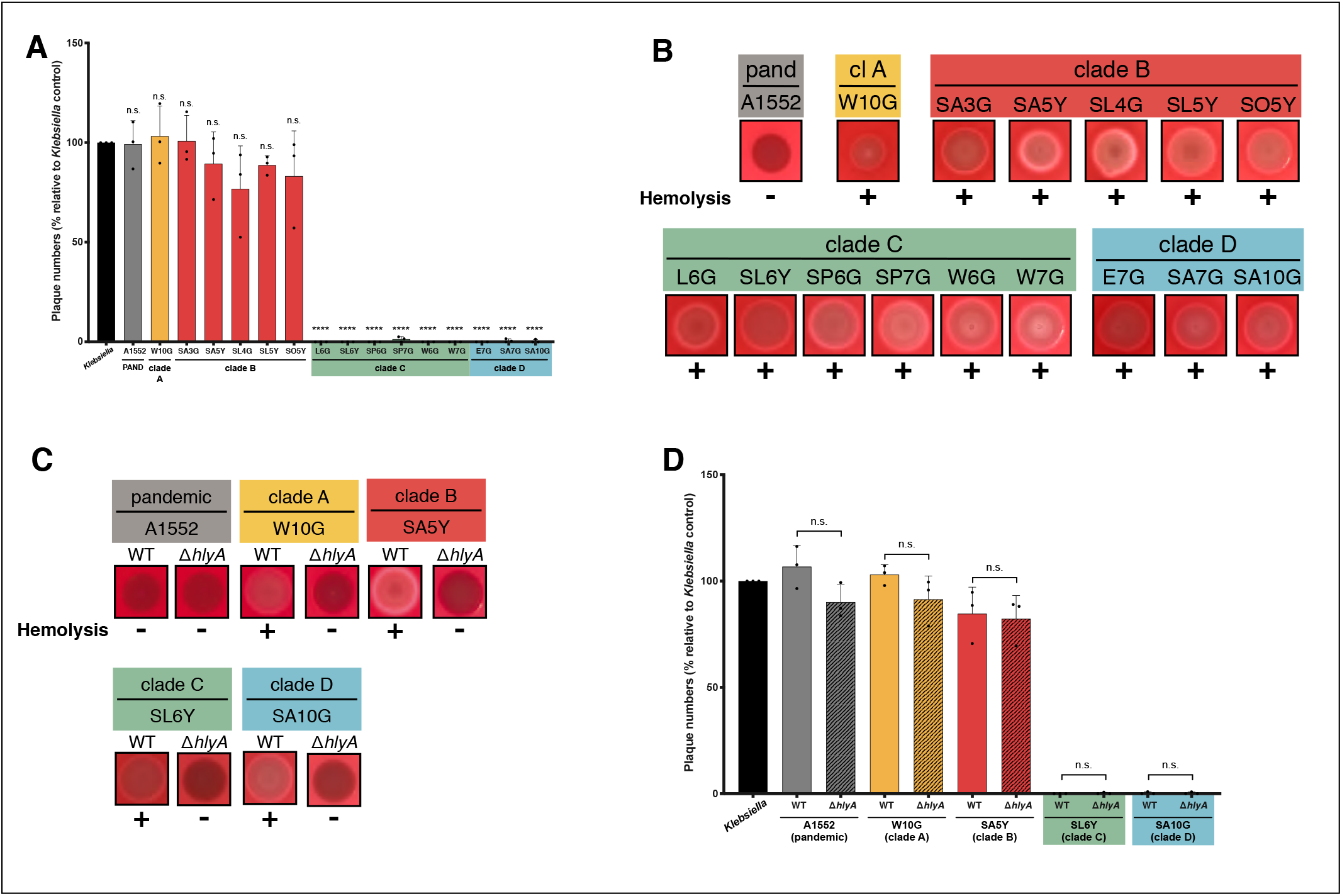
Defense against amoebal predation by a subset of *V. cholerae* strains. (A, D) Amoebal predation was scored using *D. discoideum* grazing assays in which formed plaques on bacterial lawns were enumerated. Plaque numbers are indicated relative to those formed on a lawn of *K. pneumoniae*, which served as a positive control. Bar plots represent the average of at least three independent biological replicates (±SD). Statistical significance is indicated (n.s., not significant; **** *p* < 0.0001). (B and C) Hemolytic activity was tested on blood agar plates. Pandemic *V. cholerae* strain A1552 as well as all environmental isolates (B) or a representative subset together with their respective *hlyA*-minus derivatives (C) were assessed for hemolysis.

### Enhanced hemolysin production in environmental *V. cholerae* does not contribute to their anti-amoebal behavior

As we observed that a subset of the environmental strains had a strong anti-amoebal effect, we wondered whether this grazing inhibition was linked to toxic effectors of *V. cholerae*. Our group had previously demonstrated that the proper timing of the production or activity of the pore-forming toxin hemolysin HlyA was essential for pandemic *V. cholerae* to form a replication niche inside the aquatic amoebae *A. castellanii*. Indeed, while pandemic *V. cholerae* exerted a tight regulation over this toxin and thereby successfully infected the amoebae’s contractile vacuole, constitutive hemolysin activity by the environmental isolates killed the host prematurely, preventing the formation of this replication niche (Van der Henst *et al*., 2018). We speculated that hemolysin activity could also be involved in the resistance against *D. discoideum* grazing that we observed for the environmental strains from clades C and D. To follow up on this hypothesis, we first tested all environmental WT strains for hemolysis on blood agar plates. As shown in Figure 1B, all isolates had strong hemolytic activity, especially when compared with the pandemic strain A1552. To ensure that the hemolysis was indeed caused by HlyA’s activity, we interrupted the *hlyA* gene (loci comparable to locus tag VCA0219 in reference strain N16961) in a subset of the environmental strains through the integration of an antibiotic resistance cassette (the wild-type (WT) versions of the environmental strains as well as their respective mutants are listed in Table S1). The selection of this subset of environmental isolates was thereby based on two criteria: i) to represent each clade; and ii) to select those strains that showed efficient chitin-induced natural transformability, which allowed for genetic manipulation of those strains. For these representative strains, we confirmed that the observed blood cell lysis was indeed caused by the specific hemolysin HlyA (Fig. 1C). After this confirmation, we tested the strains in the amoebal grazing assay. However, to our surprise, the hemolysin-deficient mutants behaved the same way as their parental WT strains, indicating that HlyA is not responsible for the strong anti-amoebal behavior that we observed for the clade C and D strains (Fig. 1D). These results therefore suggest that, at least under the tested conditions, HlyA does not play an important role in anti-amoebal grazing defense.

### Environmental isolates constitutively produce T6SS

As HlyA activity was ruled out as a defense mechanism, we moved on to check the potential involvement of other molecular weapons. We next considered the T6SS, due to its widespread occurrence in Gram-negative bacteria. Notably, *V. cholerae’s* T6SS was initially discovered due to its anti-eukaryotic activity that allowed the non-pandemic *V. cholerae* strain V52 to avoid predation by *D. discoideum* (Pukatzki *et al*., 2006). Indeed, previous studies had indicated that, in contrast to the 7^th^ pandemic strains, non-pandemic toxigenic *V. cholerae* (such as the two O37 serogroup strains V52 and ATCC25872; Pukatzki *et al*., 2006; Basler *et al*., 2012; Van der Henst *et al*., 2018) as well as environmental isolates (Unterweger *et al*., 2012; Bernardy *et al*., 2016; Crisan and Hammer, 2020) maintain constitutive T6SS activity. This is in contrast to the silenced T6SS of pandemic strains, which are primarily induced by chitin or low c-di-GMP levels, concomitantly with natural competence and the production of extracellular enzymes, respectively (Borgeaud *et al*., 2015; Watve *et al*., 2015; Metzger and Blokesch, 2016; Metzger *et al*., 2016; Veening and Blokesch, 2017). To check whether the T6SS protects clade C and D strains, we first tested the general T6SS activity of the environmental isolates. As shown in Figure 2A, we observed that the environmental strains efficiently eradicated *Escherichia coli* prey bacteria. Only strain SA3G of clade B reproducibly killed prey with a reduced efficiency, even though residual T6SS activity was still observed when compared with the nonkilling pandemic strain A1552 (Fig. 2A). However, these data alone do not unambiguously show whether the observed prey effacement was indeed T6SS-dependent or was instead the result of any other modes of interbacterial competition, such as contact-dependent inhibition, toxin secretion, bacteriocins, etc. (Hibbing *et al*., 2010; Stubbendieck and Straight, 2016; Granato *et al*., 2019). We therefore deleted the T6SS sheath protein-encoding gene *vipA* in each of the clade-representing strains and confirmed their lost T6SS activity by scoring for Hcp secretion. As illustrated in Figure 2B, all strains were able to produce Hcp protein but only the T6SS-active parental environmental strains were able to also secrete this protein into the supernatant. These data are therefore in agreement with the idea that the strains’ T6SS is indeed constitutively active (at least under the tested conditions) and is the reason behind the observed interbacterial killing phenotype (Fig. 2A). We confirmed the latter idea by comparing the WT and *vipA*-minus derivatives’ killing ability in an interbacterial competition assay using *E. coli* as prey (Fig. 2C).

**Figure 2:**
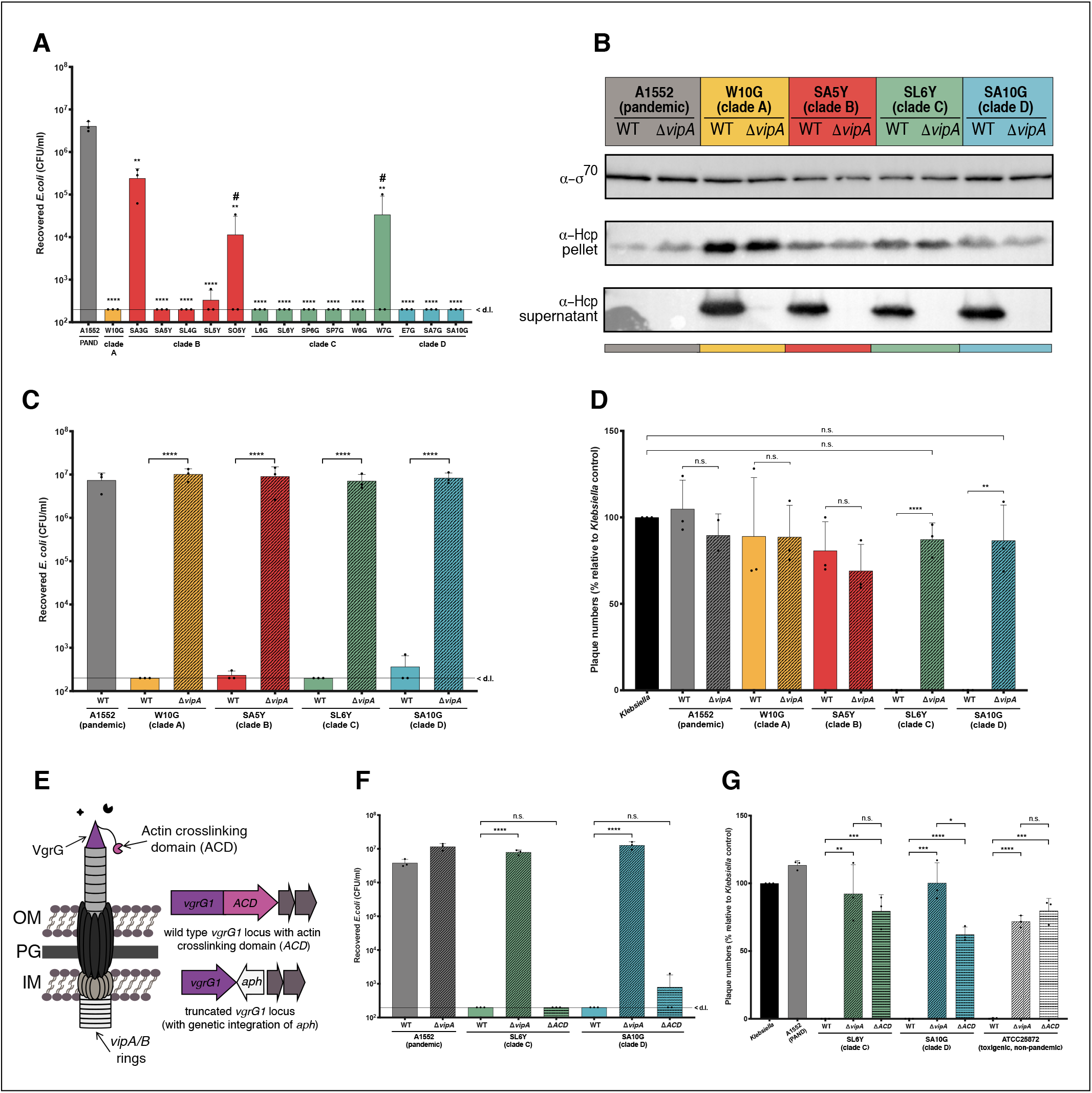
Constitutive T6SS activity linked to an ACD-containing effector inhibits amoebal grazing. (A, C, F) Bacterial killing assays using *E. coli* as prey. Numbers of surviving prey are depicted on the Y-axis (CFU/ml). Statistical significance in panel A is shown above each strains’ bar and calculated relative to the T6SS-silent negative control strain A1552. #, for these strains, the killing activity was only reduced in one of the three independent experiments. (B) T6SS activity in representative environmental strains. Hcp detection in WT and Δ*vipA* mutants of representative environmental isolates. Intracellular (pellet) and secreted (supernatant) Hcp were assessed by immunoblotting using Hcp-directed antibodies. Detection of σ70 served as a loading control. (D, G) T6SS- and ACD-dependency of the anti-amoebal defense. Plaque formation by *D. discoideum* on bacterial lawns formed by representative *V. cholerae* WT, *vipA* derivatives (D and G) and ACD-minus (G) strains. Details as in Fig. 1. The toxigenic non-pandemic strain ATCC25872 and its site-directed mutant served as control in panel G. (E) Simplified scheme of the T6SS. The actin crosslinking domain (ACD) consists of a C-terminal extension of the VgrG1 tip protein and this multidomain protein is encoded by the *vgrG1* locus (shown on the right). Removal of the ACD-encoding sequence was accomplished through site-directed integration of a stop codon concomitantly with an *aph* selective marker. Bar plots in all panels represent the average of at least three independent biological replicates (±SD). < dl, below detection limit. Statistical significance is indicated (n.s., not significant; \**p* < 0.05; ** *p* < 0.01; *** *p* < 0.001; **** *p* < 0.0001; for panel A, each sample was compared to the A1552 control).

### Environmental strains use their VgrG-linked ACD of the T6SS to fight amoebae

Having recognized that all environmental strains constitutively produce their T6SS, we moved on to assess the involvement of this machinery in the anti-amoebal defense of clade C and D strains. Indeed, the observed inhibition of amoebal plaque formation by a subset of the environmental isolates was consistent with previous work by Unterweger and colleagues (Unterweger *et al*., 2012). These authors had studied four environmental isolates from estuaries of the Rio Grande delta for anti-amoebal and anti-bacterial activity and observed that two of these isolates could not resist amoebal predation. These strains were, however, also unable to kill *E. coli* prey, and the reason for this interbacterial noncompetitiveness was a frameshift mutation in the intermediate T6SS regulatory protein-encoding gene *vasH* (Pukatzki *et al*., 2006; Unterweger *et al*., 2012). In contrast, apart from one exception (strain SA3G), all of the tested environmental isolates in our study efficiently eradicated *E. coli* prey (Fig. 2A), indicating that the T6SS was, in general, functional and active. We therefore reassessed the amoebal plaque formation against the genetically modified T6SS mutant strains. As shown in Figure 2D, clade C and D isolates, whose parental WT strains completely blocked amoebal predation, became nontoxic when their T6SS was inactivated, indicating that their anti-amoebal defense was indeed linked to the T6SS and uniquely caused by the latter.

Considering that all of the environmental isolates have a constitutively active T6SS under the tested conditions and efficiently killed other bacteria (Fig. 2A), we wondered why only the strains from clades C and D were able to use their T6SS as an anti-eukaryotic defense tool. To answer this question and to also characterize the full E/I modules of these strains, we inspected the T6SS clusters in the new genomic sequencing data and observed a clear clade specificity. Only those strains belonging to clades C and D encoded evolved VgrG1 proteins with a C-terminal actin cross-linking domain (ACD) (Figs. 2 and 3; Table S3). Strains from clades A and B, on the other hand, encoded only structurally relevant VgrG1 proteins without an evolved effector domain. Importantly, pandemic strains also encoded such an ACD as part of *vgrG1*, but, as noted above, these strains do not produce functional T6SSs without specific environmental cues and therefore show neither anti-bacterial (Fig. 2A) nor anti-amoebal behavior (Fig. 2D) under the tested conditions.

**Figure 3:**
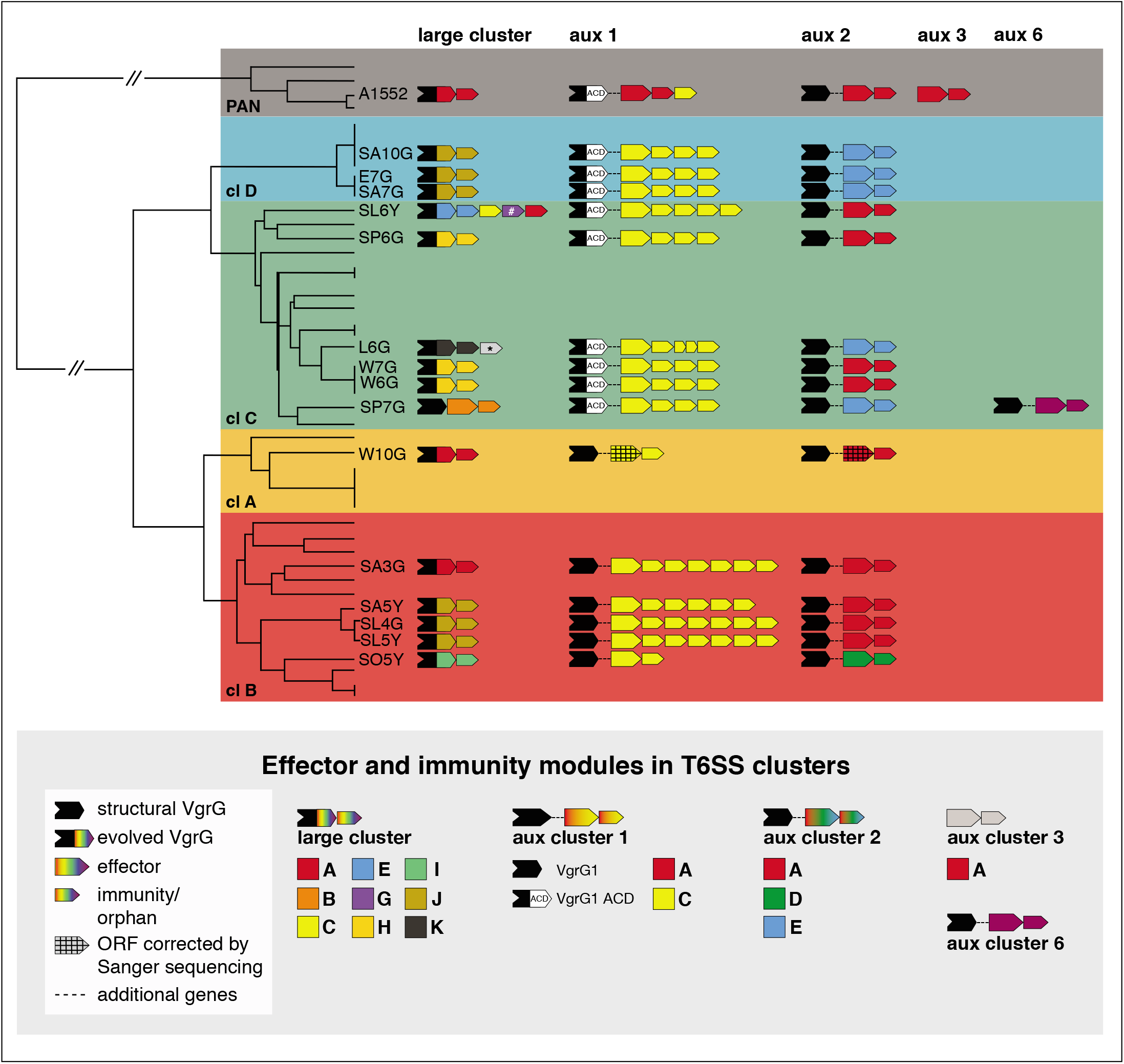
T6SS effector/immunity typing scheme of pandemic and environmental *V. cholerae*. The previously published phylogenetic tree was freely adapted from Keymer *et al*. 2007 (not fully to scale). The clade color code is depicted in the background. The E/I type of each T6SS cluster (large cluster as well as auxiliary (aux) clusters 1, 2, 3 and 6) are schematized for each strain. Large black arrows symbolize *vgrG* genes, which were classified as structural (black) or evolved (colored tips, to represent the different types of C-terminal effector domains). Large and small colored arrows represent effector and immunity genes, respectively, according to the color code indicated in the legend below the scheme. Immunity genes that are not adjacent to a putative effector gene are considered as orphan immunity loci. * depicts an orphan immunity gene from strain L6G whose gene product was slightly below the amino acid identity threshold of 30% (20.3%) relative to C-type immunity proteins. # depicts an orphan immunity gene that is a homolog to *bona fide* immunity genes in other genomes. Effector genes in auxiliary clusters 1 and 2 of strain W10G (marked in figure with a pattern) were wrongly annotated in the PacBio genome sequence due to a frameshift sequencing artifact splitting each gene into two ORFs; this sequencing error was corrected by Sanger sequencing and the corrected single gene is shown in this figure.

The ACD of VgrG1 in non-pandemic but toxigenic O37 serogroup strain V52 (Chun *et al*., 2009), which produces its T6SS constitutively, was previously shown to be involved in *V. cholerae*’s toxicity towards *D. discoideum* and macrophages (Pukatzki *et al*., 2006, 2007; Ma *et al*., 2009). Moreover, this VgrG1-ACD was also responsible for intestinal inflammation and cholera toxin-independent fluid accumulation in an infant mouse model of infection (Ma and Mekalanos, 2010). Furthermore, the VgrG1-ACD of the 7^th^ pandemic *V. cholerae* strain C6706 was implicated in alternating intestinal peristalsis of zebrafish larvae, leading to the expulsion of preinoculated commensal bacteria (Logan *et al*., 2018). However, this effect was only observable upon constitutive T6SS expression using a genetically engineered derivative of this pandemic strain in which T6SS production occurred based on artificial expression of the gene encoding the quorum sensing- and chitin-linked transcription factor QstR (Lo Scrudato and Blokesch, 2013; Borgeaud *et al*., 2015; Watve *et al*., 2015; Jaskólska *et al*., 2018; Logan *et al*., 2018).

Given this previous work on toxigenic strains, we tested whether the ACD of the clade C and D environmental isolates was likewise causative of the observed anti-amoebal response. To do so, we first generated truncated versions of VgrG1 that lacked the evolved ACD domain-containing C-terminus (Fig. 2E). Importantly, these vgrG1ΔACD strains maintained their full anti-bacterial competitiveness (Fig. 2F), which indicates that the ACD deletion did not impact the general assembly and/or activity of the T6SS machinery. However, as shown in Figure 2G, the amoebal grazing ability was restored on the lawns formed by the two vgrG1ΔACD environmental *V. cholerae* strains and resulted in equal (e.g., for strain SL6Y) or intermediate (e.g., strain SA10G) amoebal plaque numbers compared with those numbers observed for complete T6SS-defective mutants. The non-pandemic but toxigenic control strain ATCC25872 (Table S1; Aldova *et al*., 1968), which, like strain V52, is constitutive T6SS active (Van der Henst *et al*., 2018), also lost its anti-amoebal activity in the absence of the T6SS core structure or when the ACD of VgrG1 was missing (Fig. 2G). We therefore concluded that some environmental *V. cholerae* might have evolved ACD-extended VgrG-encoding genes, as the ACD’s anti-eukaryotic activity protects them from environmental grazers. Whether pandemic strains subsequently horizontally acquired the ACD encoding region as previously suggested (Kirchberger *et al*., 2017), or whether the last common ancestor between the pandemic and environmental lineages already contained this specialty, that was later then lost from some wild strains, is currently unclear.

### Diversity of T6SS effectors in environmental *V. cholerae* isolates

As briefly mentioned in the previous section, the ACD-minus mutant of the clade D strain SA10G showed reduced but still significant residual T6SS-dependent anti-amoebal activity. We therefore wondered whether this could be explained by the presence of another anti-eukaryotic effector in one or several of the T6SS clusters. Furthermore, we were also interested in characterizing the full set of E/I modules in these strains, which would allow us to speculate about the strains’ competitive potential against one another. Using the newly assembled genomic data together with the previously reported genome sequence of clade B strain SA5Y (Matthey *et al*., 2018; Matthey *et al*., 2019) we determined the E/I modules of these 15 environmental strains and predicted their function based on BlastP analyses (Fig. 3 and Table S3). Moreover, to arrange the E/I pairs into putative compatible groups, we defined their modules based on a percentage amino-acid identity of at least 30%, which is a typing approach previously applied (Unterweger *et al*., 2014; Kirchberger *et al*., 2017). Interestingly, we found evidence for several orphan immunity genes, meaning immunity genes that no longer coexisted with and were adjacent to a cognate effector-encoding gene, as had been previously reported by Kirchberger and colleagues (Kirchberger *et al*., 2017). These orphan genes were located in the 3’ regions of the T6SS clusters and were likewise classified according to their module type (Fig. 3 and Table S3). The characterization of the E/I modules and orphan immunity loci was restricted to the well-characterized T6SS clusters of *V. cholerae* (e.g., the large cluster as well as auxiliary clusters 1 and 2), as the reported auxiliary cluster 3 (E/I pair TseH/TseI; Altindis *et al*., 2015) was absent from all environmental isolates. This finding is consistent with a preprinted study showing that this auxiliary cluster represents a pandemic strain-associated mobile genetic element (Santoriello *et al*., 2019) (Fig. 3 and Table S3). Moreover, the recently characterized auxiliary clusters 4 and 5 were also absent from the environmental isolates (Labbate *et al*., 2016; Crisan *et al*., 2019). Instead, automatic annotations linked to a manual analysis identified the presence of a novel T6SS cluster in strain SP7G (clade C), which we defined as auxiliary cluster 6 (Fig. 3, Fig. S1, and Table S3). This cluster resides in the small chromosome 2 of strain SP7G and has the same gene order as auxiliary clusters 1 and 2, namely: *hcp*, *vgrG*, a putative adaptor-protein encoding gene (coding for a DUF4123 domain as described for Tap1 and VasW; Liang *et al*., 2015; Unterweger *et al*., 2015) plus two genes encoding proteins of unknown function, which we speculate represent an E/I pair. However, BlastP analysis identified no significant homologous proteins. Interestingly, our search for aux 6 cluster in the PATRIC nucleotide sequence database (Wattam *et al*., 2017) suggested that this cluster is prominently represented in several *V. cholerae* strains, including 2013 environmental isolates from Bangladesh and an O35 strain (1311-69) isolated in 1969 from a patient in India (Bishop-Lilly *et al*., 2014).

In the large T6SS cluster, pandemic *V. cholerae* strains, such as A1552 carry an A-type E/I module (Fig. 3 and Table S3), for which, in fact, the peptidoglycan cell wall degradation effector corresponds to the C-terminal domain of the evolved VgrG3 protein (Zheng *et al*., 2011; Brooks *et al*., 2013). When analyzing this locus in the environmental *V. cholerae* isolates, we noticed that the evolved nature of VgrG3 was conserved among these strains, except for clade C strain SP7G in which VgrG3 is solely a structural T6SS component that is followed by a B-type E/I module. How this effector is attached to the VgrG tip protein is, however, unclear, as no adaptor protein such as those encoded by *tap1* and *vasW* could be identified. This B-type effector is predicted to have a cellular adhesion function (Unterweger *et al*., 2014). Interestingly, strains W10G (clade A) and SA3G (clade B) contain a pandemic-like A-type E/I module (Fig. 3), while the other strains carried a wide variety of E/I pairs, as described in detail in Table S3. Moreover, even though the amino acid identity of these effectors is below 30% when comparing different types, most of them have a common predicted function, namely, peptidoglycan degradation (Table S3). Therefore, with the exception of strain SP7G, all the environmental isolates have a dedicated anti-bacterial E/I module in the T6SS large cluster (Fig. 3 and Table S3). Interestingly, pairwise comparisons of the effector and immunity protein sequences among different strains showed that strains harboring E/I modules from the same family had 100% immunity identity (Table S4). Given the diversity of E/I modules in this cluster when comparing all strains, the complete identity of immunity proteins from the same family could indicate recent acquisition by horizontal gene transfer. Furthermore, L6G and SL6Y (clade C) are the only strains that also harbor one or several orphan immunity loci after the E/I module in this large T6SS cluster (Fig. 3).

These orphan immunity loci are distinct in their type when compared with the current resident E/I modules. When we searched the PATRIC translated nucleotide sequence database (Wattam *et al*., 2017) using the protein sequence from these orphan loci as the query, we noticed that these genes are only found as orphan loci in other genomes. The only exception was the second orphan locus in strain SL6Y, which encodes a G-type immunity protein (marked by “#” in Fig. 3). The predicted protein showed 98.4% identity to true immunity proteins (e.g., those encoded directly adjacent to an effector gene) from several *V. cholerae* strains. Among those was strain 2633-78, an O1 CTX-negative isolate collected from sewage in Brazil in 1978. Interestingly, this strain was experimentally tested in a previous study, where it was shown to have an active T6SS (Bernardy *et al*., 2016).

The auxiliary cluster 1 contains the structural or evolved (e.g., encoding C-terminal ACD) *vgrG1* gene, as mentioned above (Figs. 2 and 3). In addition, in the case of pandemic *V. cholerae*, this cluster harbors an A-type E/I module encoding the lipase effector TseL with anti-bacterial and anti-eukaryotic activity (Zheng *et al*., 2011; Dong *et al*., 2013; Russell *et al*., 2013) followed by a C-type orphan immunity gene (Kirchberger *et al*., 2017). Interestingly, all of the 15 environmental strains harbor C-type E/I modules as part of this auxiliary cluster 1, even though there is considerable polymorphism in the effector and immunity proteins (Fig. 3 and Table S5). C-type effectors have a predicted alpha/beta hydrolase domain (DUF2235), which has been previously associated with T6SS phospholipases from *E. coli* and *Pseudomonas aeruginosa* (Russell *et al*., 2013; Flaugnatti *et al*., 2016; Crisan *et al*., 2019). Furthermore, with the exception of strains W10G (clade A) and SO5Y (clade B), all environmental isolates also carry between two and five orphan immunity loci downstream of the E/I pair (Fig. 3). Considering the wide variety of E/I types in the large cluster as well as in auxiliary cluster 2 (see below), the apparent conservation of C-type E/I modules in the auxiliary cluster 1 of these strains, in addition to the many C-type orphan loci (as also observed in the *in silico* study by Kirchberger *et al*. 2017) is quite intriguing and supports the idea that the C-type orphan immunity gene in the pandemic strains might be a remnant of a previous C-type E/I module of the strains’ progenitor.

In pandemic *V. cholerae*, the auxiliary cluster 2 carries an A-type E/I module where the VasX effector is a pore-forming toxin (Miyata *et al*., 2011; Miyata *et* al., 2013; Russell *et al*., 2014). Our comparative genomic analyses showed that several of the environmental isolates likewise encode VasX-like effectors (A-type) at the same locus (Fig. 3 and Tables S3 and S6), while other strains carry D- and E-type E/I modules. While D-type effectors have been predicted to foster peptidoglycan degradation, E-type effectors are predicted to form pores, like VasX (Unterweger *et al*., 2014). We therefore reasoned that the residual ACD-independent T6SS-dependent anti-amoebal impact observed for strain SA10G (clade D) might be caused by this E-type effector from auxiliary cluster 2, especially as the latter is missing from clade C strain SL6Y in which removal of the ACD from VgrG1 was almost equivalent to a complete T6SS inactivation (Fig. 2G). Interestingly, our analyses suggest that these E-type effectors also contain a common peptidoglycan-binding domain (e.g., an N-terminal Lysin Motif; Buist *et al*., 2008), which might render them bifunctional against bacteria and eukaryotes (Fig. 3 and Table S3). Future work will address the exact characteristics of the putative E/I modules in more detail.

### Competition among environmental strains occurs in the presence of nonidentical E/I modules

Previous studies showed that a plethora of T6SS-transported effectors have active anti-bacterial purposes and that cognate immunity proteins are required to protect the producer or its siblings from intoxication (Dong *et al*., 2013; Durand *et al*., 2014; Russell *et al*., 2014; Unterweger *et al*., 2014). Strains with matching E/I modules could therefore coexist in the same environment (Unterweger *et al*., 2014), while competitive strains might clear a niche and propagate inside this niche in a clonal manner (McNally *et al*., 2017; Speare *et al*., 2018). Importantly, as these E/I modules seem to move horizontally in an as yet uncharacterized manner (Unterweger *et al*., 2014; Salomon *et al*., 2015), the level of compatibility between strains will not follow their phylogenetic relatedness. We therefore sought to experimentally probe the strains’ compatibility or competitiveness within this collection of environmental isolates. Indeed, while we scored T6SS activity against a laboratory strain of *E. coli* (see Fig. 2), we wondered how the *V. cholerae* strains would behave when exposed to each other. Consistent with the fact that the E/I modules carried by the environmental strains are considerably different from those of pandemic *V. cholerae* (Fig. 3 and Table S3), we experimentally demonstrated that these bacteria efficiently outcompeted the T6SS-silent pandemic strain A1552 (Fig. 4A). Next, we tested interbacterial competition between the clade-representative strains in an assay in which the prey strains had their T6SS inactivated. As shown in Figure 4B, reciprocal killing of the T6SS-positive parental strains occurred and only kin strains were immune to the toxic assaults. However, as the E/I module types seemed more similar within clades than across clades, we extended our analyses and tested all pairwise combinations (this time, with both partners T6SS positive). The underlying rationale was that previous studies had identified different types of E/I modules *in silico* (Unterweger *et al*., 2014; Kirchberger *et al*., 2017; Crisan *et al*., 2019) but most pairwise competition assays were primarily linked to patient isolates and not to a larger collection of environmental isolates. Indeed, as described above and shown in Figure 3, our *in silico* prediction showed that strains within clades often contained similar E/I module types (Tables S4 to S6), even though the amino acid identity threshold for this categorization is rather low (30% as previously defined; Unterweger *et al*., 2014; Kirchberger *et al*., 2017).

**Figure 4:**
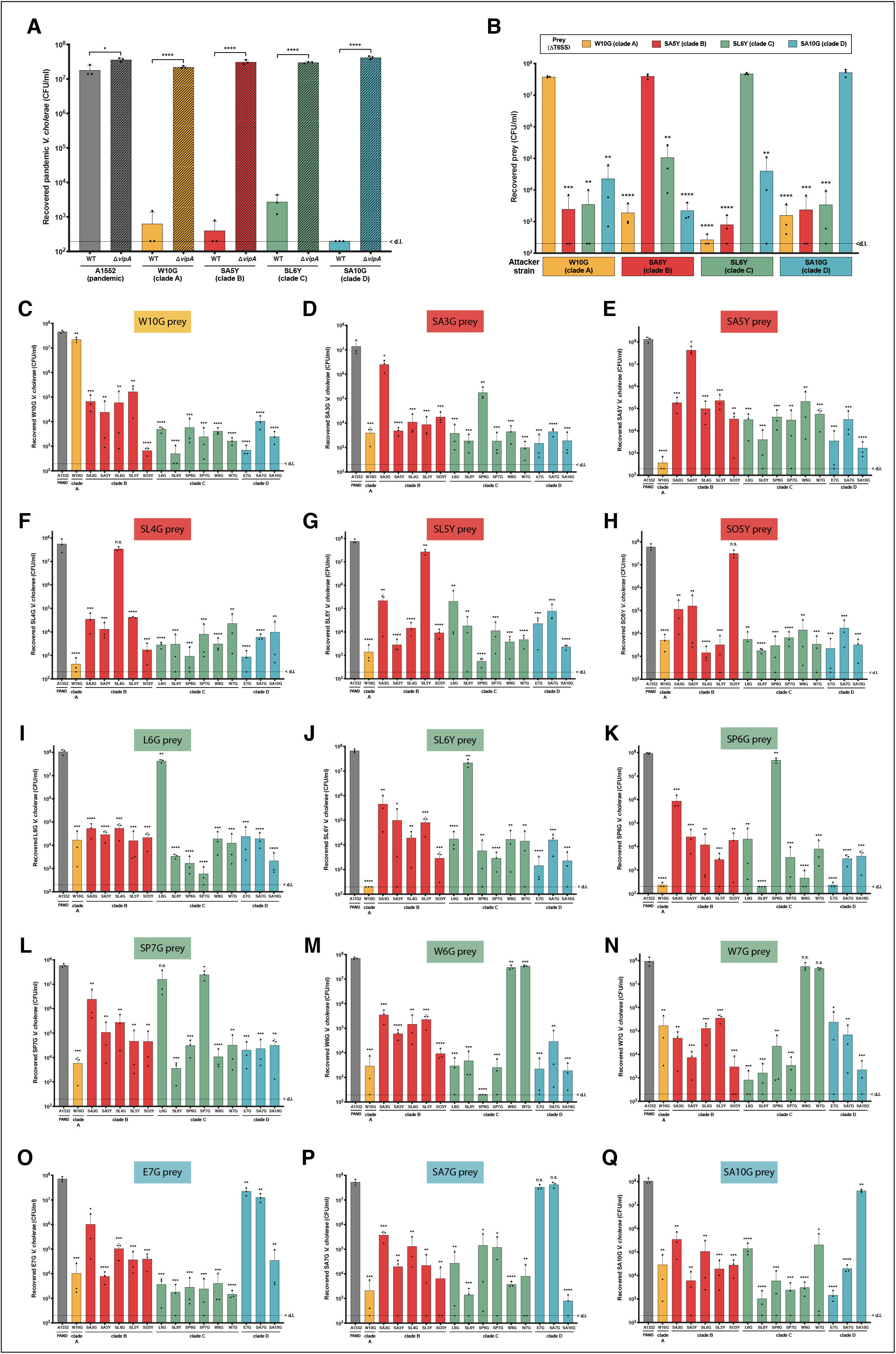
Competitiveness among environmental strains. Graphs in all panels represent bacterial killing assays as described in Fig. 2 using the predator strains as indicated on the X-axes and as prey: (A) the pandemic *V. cholerae* strain A1552; (B) representative *vipA*-minus (ΔT6SS) mutants of the environmental isolates as shown above the graph; or (C to Q) each environmental isolate as shown in the graph title for each panel and on the Y-axis. Plots represent the average of three independent biological replicates (±SD). < dl, below the detection limit. Statistical significance is indicated (n.s., not significant; * *p* < 0.05; ** *p* < 0.01; \*\*\**p* < 0.001; **** *p* < 0.0001).

As shown in Figure 4, we observed interesting and complex phenotypes in these extensive pairwise killing experiments. For instance, strains W6G and W7G from clade C were fully protected against each other’s attacks (Fig. 4 panels M and N). This finding is consistent with the 100% identity of all three T6SS clusters, including the orphan genes from aux cluster 1 (Tables S4 to S6), and their overall near clonality as described above. Clade D strains have the same E/I module types in all T6SS clusters. However, upon closer inspection of their E/I protein sequences, we can observe a 100% sequence identity among the three strains only in the large and aux 1 cluster-encoded proteins. For the aux 2 cluster, the encoded E/I proteins from strains E7G and SA7G are 100% identical, while the E/I pair carried by strain SA10G has only 95%/73% identity when compared with the other two strains’ E/I pair (Tables S4 to S6). Consequently, and as shown in Figure 4 (panel O to Q), strains E7G and SA7G are fully compatible with each other, while strain SA10G can kill and be killed by the other two strains. We therefore concluded that the identity level in only one of the T6SS cluster-encoded immunity proteins causes the competitiveness among these strains.

Surprisingly, clade B strains SA5Y and SL4G have 100% identity in all three T6SS cluster-encoded immunity proteins (Tables S4 to S6) but are still able to kill each other with considerable efficiency (Fig. 4, panels E and F). Why this is the case is currently unclear. However, we speculate that expression or immunity protein production might be impaired in those strains or that additional T6SS E/I modules are hidden in the strains’ genomes, which were not easily identifiable based on a lack of the hallmark genes *paar, hcp* and *vgrG* in their vicinity.

Clade A strain W10G carries pandemic-like A-type E/I modules in the large and aux2 clusters (Fig. 3). Interestingly, this strain can kill many of the other environmental strains remarkably well, such as clade B strains SA5Y, SL4G and SL5Y, and clade C strains SL6Y and SP6G. Notably, the T6SS active toxigenic strain V52 (which harbors A-type E/I modules in all T6SS clusters) was previously shown to outcompete strains carrying different combinations of E/I modules (Unterweger *et al*., 2014). Indeed, the AAA (A-type in all clusters) E/I modules is conserved in pandemic *V. cholerae*, even in those strains that caused former pandemics (e.g., 6^th^ pandemic O1 classical strains), as well as in non-pandemic but toxigenic isolates. It was therefore speculated that this combination might be advantageous in a disease context (Unterweger *et al*., 2014). Clade B strain SA3G also contains A-type E/I modules in the large and aux 2 clusters, but it does not kill other strains at the same level as strain W10G. However, as noted above, this strain is even less efficient against laboratory *E. coli* prey strains (Fig. 2A), which could mask its full effector toxicity potential.

Finally, a very interesting pairwise comparison is that of clade C strains SP7G and L6G. While L6G as a prey is very efficiently eliminated by strain SP7G (Fig. 4, panel I), SP7G prey is resistant to L6G intoxication (Fig. 4, panel L). These strains contain E/I sets of different families in the large cluster and show 78% and 99.7% identity in the immunity proteins encoded in aux cluster 1 and 2, respectively. Based on these differences, we would expect that these strains fully compete with each other; however, killing only occurs with SP7G as the predator. We therefore speculate that the large cluster-encoded K-type effector domains of strain L6G’s evolved VgrG3 as well as SP7G’s noncanonical structural VgrG3 and its adjacently encoded putative effector protein might not be functional peptidoglycan destruction enzymes and therefore not necessarily active in interbacterial competition. Why a low level of protein identity in the aux cluster 2-encoded immunity protein is sufficient to protect strain SP7G from L6G assaults is currently unclear. However, it is tempting to speculate that the *vice versa* interaction leads to L6G killing due to the additional aux cluster 6 that is carried by strain SP7G. Further work is therefore necessary in order to delve deeper into these observed phenotypes.

Overall, closer inspection of all of the pairwise killing data attests to the complexity behind the T6SS compatibility code. As mentioned previously, even though some strains might harbor the same E/I families in the T6SS clusters, pairwise comparisons of these proteins shows that quite frequently their identity is not 100% (Tables S4 to S6), which appears to be necessary to allow coexistence (also observed in Speare *et al*., 2018). Our data therefore support what Unterweger and colleagues (Unterweger *et al*., 2014) initially speculated, namely, that the compatibility between strains seems to follow the level of polymorphism of their immunity proteins and that diversity in only one cluster-encoded protein is sufficient to drive competition.

### Conclusion

In this study, we investigated two minor virulence factors, the pore forming hemolysin and the T6SS, in a set of fifteen environmental *V. cholerae* strains. We assessed the relevance of these molecular weapons as defense mechanisms against amoebal predation (hemolysin and T6SS) and in the context of bacterial warfare (T6SS only). We showed that all of these environmental isolates possess a constitutively active T6SS and are able to use the machinery as a bacterial killing device. In contrast, only a subset of these strains was able to efficiently suppress grazing by *D. discoideum* amoebae, a phenotype that was dependent on the eukaryote-specific ACD of the evolved VgrG1 T6SS effector. Careful *in silico* identification unveiled an extensive T6SS repertoire of E/I pairs and orphan immunity loci. Consistent with this finding, we observed extensive interbacterial competition under pairwise coculture conditions whereby mutual compatibility was rarely achieved. Importantly, our study also confirmed that both molecular weapons, the T6SS and the hemolysin toxin, are constitutively active in the environmental isolates in contrast to their tight regulation in the well-studied pandemic patient isolates. Future work is therefore required to decipher how this differential production pattern is achieved in pandemic versus non-pandemic strains and whether this tight regulatory control might provide specific benefits to the former strains.

## Experimental procedures

### Bacterial strains and growth conditions

The bacterial strains (*V. cholerae, E. coli* and *K. pneumoniae*) used in this study are listed in Table S1. Unless otherwise stated, all strains were grown aerobically in Lysogeny broth (LB; 10 g/L of tryptone, 5 g/L of yeast extract, 10 g/L of sodium chloride; Carl Roth) or on LB agar plates at 30°C. Half-concentrated defined artificial seawater medium (0.5×DASW) containing HEPES and vitamins (Meibom *et al*., 2005) was used for growth on chitinous surfaces for strain construction based on chitin-induced natural transformation (see below).

*D. discoideum* amoebae (strain Ax2 Ka) were cultured in HL5 medium supplemented with glucose (Formedium, UK). For amoebal grazing assays (e.g., plaque formation assays; see below), SM/5 medium (final concentrations: 2 g/L of glucose, 2 g/L of bacto peptone, 2 g/L of yeast extract, 0.2 g/L of MgSO_4_ 7H_2_O, 1.9 g/L of KH_2_PO_4_, 1 g/L of K_2_HPO_4_; pH 6.4), was mixed with 2% agar to prepare SM/5 plates (20 mL/plate) (Sussman, 1987). CaCl_2_ (50 μM)-supplemented Sörensen’s buffer (8 g /4 L of KH_2_PO_4_, 1.16 g /4 L of Na_2_HPO_4_; pH 6; Gerisch *et al*., 1967) was used as the washing and resuspension buffer for the amoebae and the bacteria that were used in the plaque formation assay.

The following antibiotics were added if required at the given concentration: kanamycin (75 μg/ml), rifampicin (100 μg/ml), streptomycin (100 μg/ml) and chloramphenicol (2.5 μg/ml).

### Genetic engineering

*V. cholerae* strains were genetically modified using chitin-induced transformation as previously described (Marvig 2010; Silva and Blokesch, 2010; Blokesch, 2012; Borgeaud and Blokesch, 2013). This method relies on natural transformation triggered by growth on chitin followed by the addition of a PCR fragment that carried the desired genetic change. To achieve higher numbers of transformants, the protocol was slightly modified. The PCR fragments were added twice (24h and 36h after bacterial inoculation on the chitin flakes) and cells were enriched in 2×YT medium (Carl Roth) before selective plating. PCR amplifications were conducted using Pwo (Roche) and GoTaq (Promega) polymerases according to the suppliers’ recommendations. Following initial screening by PCR (using bacterial cells as the templates), genetically engineered loci were verified by Sanger sequencing (Microsynth, Switzerland).

The rifampicin-sensitive *V. cholerae* strain A1552-Rif^S^ was generated by a combination of natural cotransformation (Dalia *et al*., 2014) and our previously described counterselectable Trans2 approach (Van der Henst *et al*., 2018). To this end, a 4kb PCR fragment was amplified, harboring a mutation, which restored the native *rpoB-encoded* protein (F531S substitution) from its mutated version in the parental strain A1552 (RpoB[S531F]; Matthey *et al*., 2018). *V. cholerae* A1552 was then cotransformed with this fragment and another 3,924 bp fragment containing flanking regions matching *lacZ* and two selection markers (*aph* and *pheS*\*; Table S1). Transformants were selected on kanamycin-containing agar plates and the lost rifampicin resistance was scored based on replica plating on plates +/- rifampicin using a velvet cloth. A second round of natural transformation followed to restore the *lacZ* gene by adding a WT *lacZ* PCR fragment to chitin-grown cells followed by a counter selection of the *pheS\** allele on 4-chloro-phenylalanine (20mM)- containing agar plates as previously described (Van der Henst *et al*., 2018). To confirm the restoration of native *rpoB*, genomic DNA of A1552 Rif^S^ was isolated and the PCR-amplified *rpoB* gene was Sanger-sequenced.

### Amoebal grazing assay

To determine the predatory capacity of *D. discoideum* on bacterial lawns of *V. cholerae*, plaque formation was scored following a previously described protocol (Pukatzki *et al*., 2006) with minor modifications. Briefly, bacteria were cultured overnight in LB medium at 30°C and harvested by centrifugation. The cell pellet was washed and resuspended in SorC buffer (Gerisch *et al*., 1967), and then diluted with SorC to reach an optical density at 600 nm (OD_600_) of 5.5 in a final volume of 400 μL. Cultured *D. discoideum* amoebae (in HL5 with glucose; Formedium, UK) were detached from culture dishes using cell scraper (SPL Life Sciences) and collected by centrifugation (3 min 1,000 rcf), resuspended in SorC buffer, and enumerated in a KOVA counting chamber (KOVA International, USA). The amoebal concentration was adjusted to 2 × 10^4^ cells/ mL and 20 μL of this suspension (corresponding to ~400 amoebal cells) was mixed with the 400 μL of bacterial suspension. The mixture was gently spread on two parallel SM/5 plates using a plastic rake (VWR), resulting in technical replicates. The plates were wrapped in aluminum foil and incubated at 24°C for 5 days. After this incubation period, *D. discoideum* plaque numbers were enumerated. As a positive control, we included a frequently used nonencapsulated *Klebsiella* strain (Benghezal *et al*., 2006) for which the resulting plaque numbers were set to 100%. Three biologically independent experiments were performed. The individual experimental data points (mean of technical replicates) as well as the overall average of the independent experiments (+/- standard deviation) are shown in each graph. A two-tailed Student’s *t*-test was performed to determine statistical significance.

### Hemolysin activity

The hemolytic activity of *V. cholerae* was assayed using trypticase soy agar containing 5% sheep blood (BD, Heidelberg, Germany). To do so, the respective overnight cultures were spotted (2 μL) onto the plates and incubated at 30°C for 24 h, after which pictures of the plates were taken.

### Interbacterial killing assays

Bacterial killing was assessed following a previously established assay with minor modifications (Borgeaud *et al*., 2015). The prey cells (*E. coli* or *V. cholerae*, as indicated) and the respective predator bacteria were mixed at a ratio of 1:10 and spotted onto paper filters on prewarmed LB agar plates. After 4 h of incubation at 37°C, the bacteria were resuspended, serially diluted, and spotted onto antibiotic-containing (rifampicin or streptomycin) LB agar plates to enumerate the colony-forming units (shown as CFU/ml). The majority of these killing experiments were performed using exponentially growing *V. cholerae* (OD_600_ ~1). For the pairwise killing experiments of all environmental strains, OD_600_-adjusted overnight cultures were used, which resulted in biologically similar outcomes to the samples derived from exponentially growing cultures. Statistically significant differences were determined on log-transformed data (Keene, 1995) by a two-tailed Student’s *t*-test of three biologically independent replicates. If no prey bacteria were recovered, the value was set to the detection limit to allow for statistical analysis.

### SDS-PAGE and western blotting

To check the production of the Hcp protein, cell lysates were prepared as described previously (Metzger *et al*., 2016). In brief, exponentially growing bacteria (~3 hours of growth after a 1:100 back dilution from overnight cultures) were pelleted and then resuspended in Laemmli buffer, adjusting for the total number of bacteria according to the cultures’ OD_600_ values. To check for T6SS-secreted Hcp, 1.5 ml of the culture supernatant was filter sterilized (0.2-μm filter; VWR) and the proteins were precipitated using trichloroacetic acid (TCA). The precipitated proteins were washed with acetone before being resuspended in 30 μL of Laemmli buffer. All samples were heated at 95°C for 15 min.

Proteins were separated by sodium dodecyl sulfate (SDS)-polyacrylamide gel electrophoresis (PAGE) using 15% gels and then western blotted as previously described (Lo Scrudato and Blokesch, 2012). Primary antibodies against Hcp (Eurogentec; Metzger *et al*., 2016) were used at 1:5,000 dilution while anti-Sigma70-HRP antibodies (BioLegend, USA distributed via Brunschwig, Switzerland) were diluted 1:10,000 and served as a loading control. Goat anti-rabbit horseradish peroxidase (HRP) (diluted at 1:20,000; Sigma-Aldrich, Switzerland) was used as the secondary antibody against the anti-Hcp primary antibody. Lumi-Light^PLUS^ western blotting substrate (Roche, Switzerland) served as the HRP substrate. The signals were detected using a ChemiDoc XRS+ station (BioRad).

### Preparation of genomic DNA for whole-genome sequencing

Genomic DNA (gDNA) was purified from 2 ml of an overnight culture of the respective strain. DNA extraction was performed using 100/G Genomic-tips together with a Genomic DNA buffer set as described in the manufacturer’s instructions (Qiagen). After precipitation, the DNA samples were washed twice with cold 70% ethanol and dissolved in Tris buffer (10 mM Tris-HCl, pH 8.0).

### Long-read PacBio genome sequencing

Sample preparation and genome sequencing was performed by the Genomic Technology Facility of the University of Lausanne (Switzerland) using standard protocols. Briefly, DNA samples were sheared in Covaris g-TUBEs to obtain fragments with a mean length of 20 kb. The sheared DNA was used to prepare each library with the PacBio SMRTbell template prep kit 1 (Pacific Biosciences) according to the manufacturer’s recommendations. The resulting library was size selected on a BluePippin system (Sage Science, Inc.) for molecules larger than 15 kb, which excluded smaller plasmids. Each library was sequenced on one single-molecule real-time (SMRT) cell with P6/C4 chemistry and MagBeads on a PacBio RS II system at a movie length of 360 min. Genome assembly was performed using the protocol RS_HGAP_Assembly.3 in SMRT Pipe 2.3.0, and circularization of the genomes was achieved using the Minimus assembler of the AMOS software package 3.1.0 using default parameters (Sommer *et al*., 2007). The assembled genomes were initially annotated using Prokka 1.12 (Seemann, 2014) but due to several incompatibilities with the NCBI database, they were reannotated with their own pipeline (PGAP annotation) during NCBI submission. The genomic data and NCBI accession numbers are summarized in Table S2.

Sequencing artifacts were observed for two genomic regions of strain W10G (clade A), leading to frameshifted putative effector genes. Briefly, for aux cluster 1, the effector gene was split into two ORFs (locus tags VC-W10G_01483 and VC-W10G_01482 in CP053794), due to the insertion of a C in position 321 of the first ORF, which led to a frameshift and consequently to an early stop codon at position 351. Similarly, in aux cluster 2, the putative effector gene was also split into two ORFs (locus tags W10G_02844 and W10G_02845 in CP053795) due to the insertion of a C in position 890, which likewise caused a frameshift followed by early termination. Both of these regions were Sanger sequenced after PCR-amplification using the same genomic DNA samples as templates that were initially used for PacBio library preparation, which confirmed that the additional C bases were in both cases a sequencing artifact and the genes were properly maintained in strain W10G (as indicated in Fig. 3).

### Characterization of E/I modules from environmental strains

T6SS clusters of the environmental strains were identified by searching the Prokka-annotated genomes for conserved genes, such as *paar, vgrG* and *hcp*, as well as according to their location when aligned to the genome of the pandemic strain A1552 (Matthey *et al*., 2018). All identified putative effectors were compared by BLAST against the NCBI database to identify conserved domains. Additional characterization was made based on the previous literature (Unterweger *et al*., 2014; Kirchberger *et al*., 2017). Furthermore, all E/I modules and orphan immunity loci were classified for their family type. Proteins with a sequence of less than 30% identity were considered as distinct incompatible types, as previously described (Unterweger *et al*., 2014; Kirchberger *et al*., 2017). For VgrG3 proteins, the typing was only based on the effector portion of the protein (Unterweger *et al*., 2014). To determine this part of the protein, the full VgrG3 sequence from all strains was aligned. The conserved region corresponding to the VgrG part of the protein was subsequently removed, which left only the variable C-terminal effector domain. This part was then used for typing and pairwise comparisons. The putative orphan immunity proteins were queried against a translated nucleotide database (PATRIC; Wattam *et al*., 2017) to identify homologous *bona fide* immunity proteins encoded adjacent to an effector gene in the other T6SS clusters.

## Data availability

PacBio raw reads of the 14 whole-genome sequenced strains have been deposited in NCBI’s Sequence Read Archive (SRA) under Bioproject accession number PRJNA633476. Details on the SRA accession numbers, BioSamples, and individual accession numbers of the *de novo* assembled and circularized genomes are provided in Table S2.

## Acknowledgments

The authors acknowledge Sandrine Stutzmann for technical assistance, A. Boehm for sharing the published environmental *V. cholerae* isolates, Lisa Metzger for provision of strain constructs, and Thierry Soldati for sharing the *D. discoideum* amoebae. We also thank Frédérique Le Roux and current and former members of the Blokesch group and of the Lausanne/Geneva amoeba club for fruitful discussions. The authors acknowledge the staff of the Lausanne Genomic Technologies Facility at the University of Lausanne for sample processing, sequencing, and genome assembly, the NCBI submission staff for help with data deposition, and Anne-Catherine Portmann and Ivan Mateus for preliminary analysis of the genome data. We also note that the purchase of the Pacific Biosciences RS II instrument at the University of Lausanne was partially supported by the Loterie Romande through the Fondation pour la Recherche en Médecine Génétique. This work was supported by the Swiss National Science Foundation (310030_185022), the Novartis Foundation for medical-biological Research (#18C178), and a consolidator grant by the European Research Council (724630). MB is a Howard Hughes Medical Institute (HHMI) International Research Scholar (grant 55008726).

## Supporting tables and figure legends

**Table S1.** *Vibrio cholerae, Escherichia coli* and *Klebsiella pneumoniae* strains used in this study.

**Table S2.** Information of the long-read whole genome sequencing data and assemblies of NCBI BioProject PRJNA633476.

**Table S3.** BlastP-predicted T6SS effector and immunity proteins^1^ of the environmental *V. cholerae* isolates.

**Table S4.** Matrices of % identity (percentage of residues that are identical) among the effector and immunity proteins harbored in the T6SS large cluster from the *V. cholerae* environmental strains and the pandemic A1552 strain.

**Table S5.** Matrices of % identity (percentage of residues that are identical) among the effector and immunity proteins harbored in the T6SS auxiliary cluster 1 from the *V. cholerae* environmental strains and the pandemic A1552 strain.

**Table S6.** Matrices of % identity (percentage of residues that are identical) among the effector and immunity proteins harbored in the T6SS auxiliary cluster 2 from the *V. cholerae* environmental strains and the pandemic A1552 strain.

**Supplementary Fig. S1: Scheme of T6SS auxiliary cluster 6 of strain SP7G.** The 4.1-kb long auxiliary T6SS cluster carried on the chromosome 2 of strain SP7G (locus tags VC-SP7G_03229 to VC-SP7G_03234 from CP053809) is depicted. Gene symbols are explained below the scheme.

