## Supporting Information for "Interbacterial competition and anti-predatory behavior of environmental *Vibrio cholerae* strains"

### Supporting figures and legends

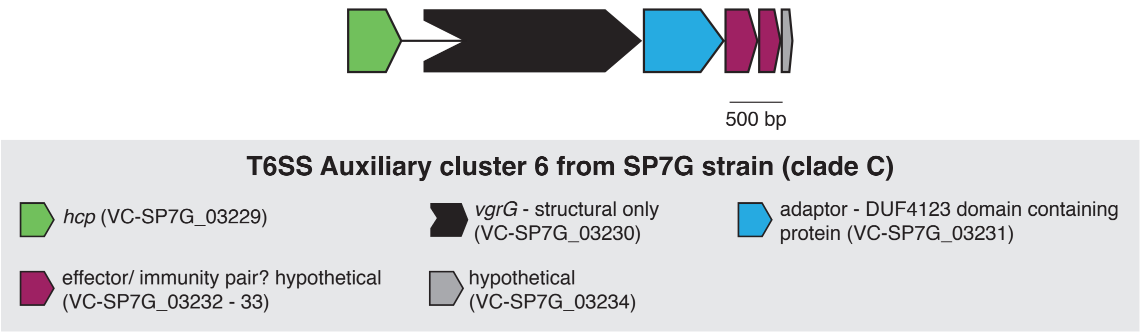

**Supplementary figure 1: Scheme of T6SS auxiliary cluster 6 of strain SP7G.** The 4.1-kb long auxiliary T6SS cluster carried on the chromosome 2 of strain SP7G (locus tags VC-SP7G_03229 to VC-SP7G_03234 from CP053809) is depicted. Gene symbols are explained below the scheme.

**Table S1.** *Vibrio cholerae*, *Escherichia coli* and *Klebsiella pneumoniae* strains used in this study.

| Strain names | Genotype / description | Internal strain number | Reference |
|---|---|---|---|
| ***V. cholerae*** | | | |
| A1552 (WT) | Wild-type, O1 El Tor Inaba; $Rif^R$ | MB_1 | (Yildiz and Schoolnik, 1998) |
| W10G | Environmental isolate (clade A) collected in Waddell Creek (CA, USA) in October 2004 | MB_5537 | (Keymer *et al.*, 2007) |
| SA3G | Environmental isolate (clade B) collected in Old Salinas River (CA, USA) in March 2004 | MB_957 | (Keymer *et al.*, 2007) |
| SA5Y | Environmental isolate (clade B) collected in Old Salinas River (CA, USA) in May 2004 | MB_353 | (Keymer *et al.*, 2007) |
| SL4G | Environmental isolate (clade B) collected in San Lorenzo River (CA, USA) in April 2004 | MB_955 | (Keymer *et al.*, 2007) |
| SL5Y | Environmental isolate (clade B) collected in San Lorenzo River (CA, USA) in May 2004 | MB_954 | (Keymer *et al.*, 2007) |
| SO5Y | Environmental isolate (clade B) collected in Soquel Creek (CA, USA) in May 2004 | MB_960 | (Keymer *et al.*, 2007) |
| L6G | Environmental isolate (clade C) collected in Lagunitas Creek (CA, USA) in June 2004 | MB_956 | (Keymer *et al.*, 2007) |
| SL6Y | Environmental isolate (clade C) collected in San Lorenzo River (CA, USA) in June 2004 | MB_953 | (Keymer *et al.*, 2007) |
| SP6G | Environmental isolate (clade C) collected in San Pedro Creek (CA, USA) in June 2004 | MB_964 | (Keymer *et al.*, 2007) |
| SP7G | Environmental isolate (clade C) collected in San Pedro Creek (CA, USA) in July 2004 | MB_952 | (Keymer *et al.*, 2007) |
| W6G | Environmental isolate (clade C) collected in Waddell Creek (CA, USA) in June 2004 | MB_354 | (Keymer *et al.*, 2007) |
| W7G | Environmental isolate (clade C) collected in Waddell Creek (CA, USA) in July 2004 | MB_962 | (Keymer *et al.*, 2007) |
| E7G | Environmental isolate (clade D) collected in Moss Landing Harbor (CA, USA) in July 2004 | MB_963 | (Keymer *et al.*, 2007) |
| SA7G | Environmental isolate (clade D) collected in Old Salinas River (CA, USA) in July 2004 | MB_959 | (Keymer *et al.*, 2007) |
| SA10G | Environmental isolate (clade D) collected in Old Salinas River (CA, USA) in October 2004 | MB_5539 | (Keymer *et al.*, 2007) |
| A1552Δ*hlyA* | A1552 with *aph* cassette in *hlyA* (A1552VC_A02993; homolog to VCA0219*); $Rif^R$, $Kan^R$ | MB_3934 | This study |
| W10GΔ*hlyA* | W10G with *aph* cassette in *hlyA* (VC-W10G_03027); $Kan^R$ | MB_8548 | This study |
| SA5YΔ*hlyA* | SA5Y with *aph* cassette in *hlyA* (VC-Sa5Y_02920); $Kan^R$ | MB_4622 | This study |
| SL6YΔ*hlyA* | SL6Y with *cat* cassette in *hlyA* (VC-SL6Y_02871); $Cm^R$ | MB_8549 | This study |
| SA10GΔ*hlyA* | SA10G with *aph* cassette in *hlyA* (VC-SA10G_02949); $Kan^R$ | MB_8550 | This study |
| A1552Δ*vipA* | A1552 with *aph* cassette in *vipA* (A1552VC_A02892; homolog to VCA0107); $Rif^R$, $Kan^R$ | MB_3012 | This study |
| W10GΔ*vipA* | W10G with *aph* cassette in *vipA* (VC-W10G_02924); $Kan^R$ | MB_6850 | This study |
| SA5YΔ*vipA* | SA5Y with *aph* cassette in *vipA* (VC-Sa5Y_02822); $Kan^R$ | MB_5772 | This study |
| SL6YΔ*vipA* | SL6Y with *aph* cassette in *vipA* (VC-SL6Y_02766); $Kan^R$ | MB_8551 | This study |
| SA10GΔ*vipA* | SA10G with *aph* cassette in *vipA* gene (VC-SA10G_02849); $Kan^R$ | MB_8552 | This study |
| A1552-$Rif^S$ | A1552 sensitive to Rifampicin (please check Material and Methods for details on construction) | MB_5161 | This study |
| A1552ΔlacZ::FRT-Kan-pheS*-FRT | A1552 with construct ΔlacZ::FRT-Kan-pheS*-FRT; $Rif^R$, $cPhe^S$ | MB_4609 | (Van der Henst *et al.*, 2018) |
| A1552-$Rif^S$Δ*vipA* | A1552-$Rif^S$ with *aph* cassette in *vipA*; $Rif^R$, $Kan^R$ | MB_8553 | This study |
| SL6Y-vgrG1ΔACD | SL6Y with truncated *vgrG1* through insertion of a stop codon and insertion of an *aph* cassette that replaces the ACD-encoding sequence; $Kan^R$ | MB_8557 | This study |
| SA10G-vgrG1ΔACD | SA10G with truncated *vgrG1* through insertion of a stop codon and insertion of an *aph* cassette that replaces the ACD-encoding sequence; $Kan^R$ | MB_8558 | This study |

| | | | |
|---|---|---|---|
| ATCC25872 | *V. cholerae* non-O1 strain (O37); isolated in 1965, Czechoslovakia; intermediate resistant to Strep | MB_276 | Aldová *et al*., 1968 |
| ATCC25872Δ*vipA* | ATCC25872 deleted for *vipA* (TransFLP); intermediate resistant to Strep | MB_5065 | Van der Henst *et al*., 2018 |
| ATCC25872 - vgrG1ΔACD | ATCC25872 with truncated *vgrG1* through insertion of a stop codon and insertion of an *aph* cassette that replaces the ACD-encoding sequence; $Kan^R$, intermediate resistant to Strep | MB_8559 | This study |
| W10G-$Rif^R$ | W10G selected for rifampicin resistance | MB_8562 | This study |
| SA3G-$Rif^R$ | SA3G selected for rifampicin resistance | MB_8563 | This study |
| SA5Y-$Rif^R$ | SA5Y selected for rifampicin resistance | MB_8564 | This study |
| SL4G-$Rif^R$ | SL4G selected for rifampicin resistance | MB_8565 | This study |
| SL5Y-$Rif^R$ | SL5Y selected for rifampicin resistance | MB_8566 | This study |
| SO5Y-$Rif^R$ | SO5Y selected for rifampicin resistance | MB_8567 | This study |
| L6G-$Rif^R$ | L6G selected for rifampicin resistance | MB_8568 | This study |
| SL6Y-$Rif^R$ | SL6Y selected for rifampicin resistance | MB_8569 | This study |
| SP6G-$Rif^R$ | SP6G selected for rifampicin resistance | MB_8570 | This study |
| SP7G-$Rif^R$ | SP7G selected for rifampicin resistance | MB_8571 | This study |
| W6G-$Rif^R$ | W6G selected for rifampicin resistance | MB_8572 | This study |
| W7G-$Rif^R$ | W7G selected for rifampicin resistance | MB_8573 | This study |
| E7G-$Rif^R$ | E7G selected for rifampicin resistance | MB_8574 | This study |
| SA7G-$Rif^R$ | SA7G selected for rifampicin resistance | MB_8575 | This study |
| SA10G-$Rif^R$ | SA10G selected for rifampicin resistance | MB_8576 | This study |
| ***E. coli and K. pneumoniae*** | | | |
| TOP 10 | F- mcrA Δ(mrr-hsdRMS-mcrBC) φ80lacZΔM15 ΔlacX74 nupG recA1 araD139 Δ(ara-leu)7697 galE15 galK16 rpsL(StrR) endA1 λ- | MB_741 | This study |
| *K. pneumoniae* | *Klebsiella* sp. / pANT5 (GFP) | MB_6380 | (Benghezal *et al*., 2006) |

*reference locus tags belong to reference strain N16861 according to Heidelberg *et al*., 2000.

**Table S2.** Information of the long-read whole genome sequencing data and assemblies of NCBI BioProject PRJNA633476.

| Strain name | Strain ID | BioSample | GenBank accession numbers | Number of bases | Number of reads | Mean read length | Total number of contigs | Contig length after circularization | Total genome size | Mean coverage | GC% content |
|---|---|---|---|---|---|---|---|---|---|---|---|
| **W10G** | MB#5537 | SAMN14944057 | CP053794 (chr1) CP053795 (chr2) | 2,050,336,308 bp | 119,162 | 17,206 bp | 2 (chr1+chr2) | 3,033,188 bp (chr1) 1,028,867 bp (chr2) | 4,062,055 bp | 292 x | 47.7% (chr1) 47.2% (chr2) |
| **SA3G** | MB#957 | SAMN14944058 | CP053744 (chr1) CP053745 (chr2) | 1,239,715,817 bp | 70,375 | 17,615 bp | 2 (chr1+chr2) | 3,020,084 bp (chr1) 1,159,698 bp (chr2) | 4,179,782 bp | 201 x | 47.6% (chr1) 47.0% (chr2) |
| **SA5Y#** | MB#353 | SAMN08813238 | CP028892 (chr1) CP028893 (chr2) | 635,540,812 bp | 35,390 | 17,958 bp | 2 (chr1+chr2) | 2,955,400 bp (chr1) 1,095,478 bp (chr2) | 4,050,878 bp | 133 x | 47.8% (chr1) 46.8% (chr2) |
| **SL4G** | MB#955 | SAMN14944065 | CP053796 (chr1) CP053797 (chr2) | 1,083,962,388 bp | 55,665 | 19,472 bp | 2 (chr1+chr2) | 2,954,462 bp (chr1) 1,173,143 bp (chr2) | 4,127,605 bp | 186 x | 47.8% (chr1) 46.4% (chr2) |
| **SL5Y** | MB#954 | SAMN14944067 | CP053798 (chr1) CP053799 (chr2) | 1,623,840,987 bp | 84,121 | 19,303 bp | 2 (chr1+chr2) | 2,947,299 bp (chr1) 1,086,276 bp (chr2) | 4,033,575 bp | 336 x | 47.8% (chr1) 46.9% (chr2) |
| **SO5Y** | MB#960 | SAMN14944069 | CP053800 (chr1) CP053801 (chr2) | 865,600,319 bp | 47,657 | 18,163 bp | 2 (chr1+chr2) | 2,947,270 bp (chr1) 1,049,609 bp (chr2) | 3,996,879 bp | 245 x | 47.9% (chr1) 47.0% (chr2 |
| **L6G** | MB#956 | SAMN14944070 | CP053802 (chr1) CP053803 (chr2) | 1,134,851,816 bp | 70,295 | 16,144 bp | 2 (chr1+chr2) | 2,829,740 bp (chr1) 1,195,206 bp (chr2) | 4,024,946 bp | 225 x | 48.1% (chr1) 46.3% (chr2) |
| **SL6Y** | MB#953 | SAMN14944072 | CP053804 (chr1) CP053805 (chr2) | 826,125,289 bp | 48,156 | 17,155 bp | 2 (chr1+chr2) | 2,900,064 bp (chr1) 1,143,503 bp (chr2) | 4,043,567 bp | 175 x | 47.8% (chr1) 46.7% (chr2) |
| **SP6G** | MB#964 | SAMN14944080 | CP053806 (chr1) CP053807 (chr2) | 874,659,808 bp | 47,923 | 18,251 bp | 2 (chr1+chr2) | 2,947,818 bp (chr1) 1,229,641 bp (chr2) | 4,177,459 bp | 154 x | 47,8% (chr1) 46.7% (chr2) |
| **SP7G** | MB#952 | SAMN14944081 | CP053808 (chr1) CP053809 (chr2) | 1,339,636,117 bp | 73,934 | 18,119 bp | 3 (chr1+chr2) | 2,868,038 bp (chr1) 1,217,051 bp (chr2) | 4,085,089 bp | 217 x | 48.0% (chr1) 46.5% (chr2) |
| **W6G** | MB#354 | SAMN14944082 | CP053810 (chr1) CP053811 (chr2) CP053812 (plasmid) | 1,247,754,152 bp | 69,891 | 17,852 bp | 3 (chr1+chr2+ plasmid) | 2,967,360 bp (chr1) 1,202,132 bp (chr2) 306,484 bp (plas) | 4,475,976 bp | 192 x | 47.8% (chr1) 46.6% (chr2) 42.2% (plas) |
| **W7G** | MB#962 | SAMN14944083 | CP053813 (chr1) | 1,490,677,399 bp | 86,303 | 17,272 bp | 3 (chr1+chr2+ plasmid) | 2,967,793 bp (chr1) 1,202,155 bp (chr2) 306,481 bp (plas) | 4,476,429 bp | 223 x | 47.8% (chr1) 46.6% (chr2) 42.2% (plas) |

| | | | | | | | | | | | |
|---|---|---|---|---|---|---|---|---|---|---|---|
| | | | CP053814 (chr2) CP053815 (plasmid) | | | | | | | | |
| **E7G** | MB#963 | SAMN14944111 | CP053822 (chr1) CP053823 (chr2) CP053824 (plasmid) | 1,477,482,932 bp | 83,856 | 17,619 bp | 3 (chr1+chr2+ plasmid) | 3,040,343 bp (chr1) 1,199,805 bp (chr2) 80,726 bp (plas) | 4,320,874 bp | 194 x | 47.8% (chr1) 46.5% (chr2) 46.3% (plas) |
| **SA7G** | MB#959 | SAMN14944112 | CP053816 (chr1) CP053817 (chr2) CP053818 (plasmid 1) CP053819 (plasmid 2) | 1,689,606,494 bp | 98,160 | 17,212 bp | 4 (chr1+chr2+ plasmid 1 + 2) | 3,004,584 bp (chr1) 1,182,929 bp (chr2) 79,397 bp (plas1) 80,721 bp (plas2) | 4,347,631 bp | 245 x | 47.8% (chr1) 46.5% (chr2) 42.2% (plas1) 46.3% (plas2) |
| **SA10G** | MB#5539 | SAMN14944113 | CP053820 (chr1) CP053821 (chr2) | 1,643,277,942 bp | 91,387 | 17,981 bp | 2 (chr1+chr2) | 3,010,605 bp (chr1) 1,166,701 bp (chr2) | 4,177,306 bp | 269 x | 47.8% (chr1) 46.6% (chr2) |

#Data based on Matthey *et al*., 2018

**Table S3.** BlastP-predicted T6SS effector and immunity proteins[1] of pandemic, non-pandemic toxigenic, and environmental *V. cholerae* strains.

| Clade | Strain name | Major cluster E/I | Aux cluster 1 *vgrG1* | Aux cluster 1 E/I | Aux cluster 2 E/I | Aux cluster 3 E/I |
|---|---|---|---|---|---|---|
| Pandemic | A1552 | Encoded protein with 1018 aa; evolved VgrG3 with predicted peptidoglycan hydrolase domain; A-type. | Encoded protein with 1164 aa; evolved VgrG1 with an actin crosslinking domain (ACD). | Encoded protein with 642 aa; lipase class 3; A-type. | Encoded protein with 1086 aa; VasX; pore forming activity[2]; A-type. | Encoded protein with 224 aa; papain-like NlpC/P60 peptidase[3]; A-type. |
| D | SA10G | Encoded protein with 995 aa; evolved VgrG3 with predicted peptidoglycan hydrolase and pesticin-like lysozyme domains; J-type. | Encoded protein with 1164 aa; evolved VgrG1 with an actin crosslinking domain (ACD). | Encoded protein with 710 aa; uncharacterized alpha/beta hydrolase domain (DUF2235); C-type. | Encoded protein with 1194 aa; predicted lysin motif (binds peptidoglycan), but also with pore formation activity[4]; E-type. | Aux cluster 3 is absent. |
| | E7G | Encoded protein with 995 aa; evolved VgrG3 with predicted peptidoglycan hydrolase and pesticin-like lysozyme domains; J-type. | Encoded protein with 1164 aa; evolved VgrG1 with an actin crosslinking domain (ACD). | Encoded protein with 710 aa; uncharacterized alpha/beta hydrolase domain (DUF2235); C-type. | Encoded protein with 1197 aa; predicted lysin motif (binds peptidoglycan), but also with pore formation activity[4]; E-type. | Aux cluster 3 is absent. |
| | SA7G | Encoded protein with 995 aa; evolved VgrG3 with predicted peptidoglycan hydrolase and pesticin-like lysozyme domains; J-type. | Encoded protein with 1164 aa; evolved VgrG1 with an actin crosslinking domain (ACD). | Encoded protein with 710 aa; uncharacterized alpha/beta hydrolase domain (DUF2235); C-type. | Encoded protein with 1197 aa; predicted lysin motif (binds peptidoglycan), but also with pore formation activity[4]; E-type. | Aux cluster 3 is absent. |
| C | SL6Y | Encoded protein with 977 aa; evolved VgrG3 with a predicted peptidoglycan hydrolase and phage-like lysozyme domains. E-type. | Encoded protein with 1164 aa; evolved VgrG1 with an actin crosslinking domain (ACD). | Encoded protein with 710 aa; uncharacterized alpha/beta hydrolase domain (DUF2235); C-type. | Encoded protein with 1086 aa; 98.3% identical to A1552's VasX; pore forming activity[2]; A-type. | Aux cluster 3 is absent. |
| | SP6G | Encoded protein with 1052 aa; evolved VgrG3 with a predicted peptidoglycan hydrolase domain; H-type. | Encoded protein with 1164 aa; evolved VgrG1 with an actin crosslinking domain (ACD). | Encoded protein with 710 aa; uncharacterized alpha/beta hydrolase domain (DUF2235); C-type. | Encoded protein with 1087 aa; 94.8% identical to A1552's VasX; pore forming activity[2]; A-type. | Aux cluster 3 is absent. |
| | L6G | Encoded protein with 989 aa; evolved VgrG3 with a predicted peptidoglycan hydrolase and lambda phage-like lysozyme domains; K-type. | Encoded protein with 1167 aa; evolved VgrG1 with an actin crosslinking domain (ACD). | Encoded protein with 710 aa; uncharacterized alpha/beta hydrolase domain (DUF2235); C-type. | Encoded protein with 1194 aa; predicted lysin motif (binds peptidoglycan), but also with pore formation activity[4]; E-type. | Aux cluster 3 is absent. |
| | W6G | Encoded protein with 1052 aa; evolved VgrG3 with a predicted peptidoglycan hydrolase domain; H-type. | Encoded protein with 1164 aa; evolved VgrG1 with an actin crosslinking domain (ACD). | Encoded protein with 710 aa; uncharacterized alpha/beta hydrolase domain (DUF2235); C-type. | Encoded protein with 1088 aa; 94.1% identical to A1552's VasX; pore forming activity[2]; A-type. | Aux cluster 3 is absent. |
| | W7G | Encoded protein with 1052 aa; evolved VgrG3 with a predicted peptidoglycan hydrolase domain; H-type. | Encoded protein with 1164 aa; evolved VgrG1 with an actin crosslinking domain (ACD). | Encoded protein with 710 aa; uncharacterized alpha/beta hydrolase domain (DUF2235); C-type. | Encoded protein with 1088 aa; 94.1% identical to A1552's VasX; pore forming activity[2]; A-type. | Aux cluster 3 is absent. |

| | | | | | | |
|---|---|---|---|---|---|---|
| | **SP7G*** | Encoded protein with 439 aa, independent from VgrG3 (structural only); cell adhesion function[4]; B-type. | Encoded protein with 1164 aa; evolved VgrG1 with an actin crosslinking domain (ACD). | Encoded protein with 710 aa; uncharacterized alpha/beta hydrolase domain (DUF2235); C-type. | Encoded protein with 1194 aa; predicted lysin motif (binds peptidoglycan), but also with pore formation activity[4]; E-type. | Aux cluster 3 is absent. |
| **A** | **W10G** | Encoded protein with 1018 aa; evolved VgrG3 with predicted peptidoglycan hydrolase domain; A-type. | Encoded protein with 684 aa; structural VgrG1 protein. | PacBio sequencing artifact in the effector gene, where the ORF was split into two genes (locus tags VC-W10G_01483 and VC-W10G_01482). Region was Sanger sequenced and manually corrected before translation to allow typing. Encoded protein with 710 aa; uncharacterized alpha/beta hydrolase domain (DUF2235); C-type. | PacBio sequencing artifact in the effector gene, where the ORF was split into two genes (locus tags W10G_02844 and W10G_02845). Region was Sanger sequenced and manually corrected before translation to allow typing. Encoded protein with 1086 aa; VasX; pore forming activity[2]; A-type. | Aux cluster 3 is absent. |
| **B** | **SA3G** | Encoded protein with 1018 aa; evolved VgrG3 with predicted peptidoglycan hydrolase domain; A-type. | Encoded protein with 684 aa; structural VgrG1 protein. | Encoded protein with 710 aa; uncharacterized alpha/beta hydrolase domain (DUF2235); C-type. | Encoded protein with 1087 aa; 94.3% identical to A1552's VasX; pore forming activity[2]; A-type. | Aux cluster 3 is absent. |
| | **SA5Y** | Encoded protein with 995 aa; evolved VgrG3 with predicted peptidoglycan hydrolase and pesticin-like lysozyme domains; J-type. | Encoded protein with 684 aa; structural VgrG1 protein. | Encoded protein with 710 aa; uncharacterized alpha/beta hydrolase domain (DUF2235); C-type. | Encoded protein with 1088 aa; 93.7% identical to A1552's VasX; pore forming activity[2]; A-type. | Aux cluster 3 is absent. |
| | **SL4G** | Encoded protein with 995 aa; evolved VgrG3 with predicted peptidoglycan hydrolase and pesticin-like lysozyme domains; J-type. | Encoded protein with 684 aa; structural VgrG1 protein. | Encoded protein with 710 aa; uncharacterized alpha/beta hydrolase domain (DUF2235); C-type. | Encoded protein with 1088 aa; 93.7% identical to A1552's VasX; pore forming activity[2]; A-type. | Aux cluster 3 is absent. |
| | **SL5Y** | Encoded protein with 995 aa; evolved VgrG3 with predicted peptidoglycan hydrolase and pesticin-like lysozyme domains; J-type. | Encoded protein with 684 aa; structural VgrG1 protein. | Encoded protein with 710 aa; uncharacterized alpha/beta hydrolase domain (DUF2235); C-type. | Encoded protein with 1088 aa; 93.7% identical to A1552's VasX; pore forming activity[2]; A-type. | Aux cluster 3 is absent. |
| | **SO5Y** | Encoded protein with 1022 aa; evolved VgrG3 with predicted peptidoglycan hydrolase and muraidase (N-acetylmuramidase) domains; I-type. | Encoded protein with 684 aa; structural VgrG1 protein. | Encoded protein with 710 aa; uncharacterized alpha/beta hydrolase domain (DUF2235); C-type. | Encoded protein with 1220 aa; predicted lysin motif (binds peptidoglycan); D-type. | Aux cluster 3 is absent. |
| **Non-pandemic** | **ATCC25872** | Encoded protein with 1018 aa; evolved VgrG3 with predicted peptidoglycan hydrolase domain; A-type. | Encoded protein with 1164 aa; evolved VgrG1 with an actin crosslinking domain (ACD). | Encoded protein with 642 aa; lipase class 3; A-type. | Encoded protein with 1086 aa; VasX; pore forming activity[2]; A-type. | Encoded protein with 224 aa; papain-like NlpC/P60 peptidase[3]; A-type. |

[1] Typing according to Unterweger *et al.*, 2014 & Kirchberger *et al.*, 2017.
[2] According to Miyata *et al.*, 2011.
[3] According to Hersch *et al.*, 2020.
[4] According to Unterweger *et al.*, 2014.
* Strain SP7G (clade C) contains an additional T6SS cluster, called Aux 6, which has the same gene order as auxiliary clusters 1 and 2 (see Fig. S1).

**Table S4.** Matrices of % identity (percentage of residues that are identical) among the effector and immunity proteins harbored in the T6SS large cluster from the *V. cholerae* environmental strains and the pandemic A1552 strain.

**Large cluster – Effector**

| | A1552 | SA10G | E7G | SA7G | SL6Y | SP6G | L6G | W7G | W6G | SP7G | W10G | SA3G | SA5Y | SL4G | SL5Y | SO5Y |
|---|---|---|---|---|---|---|---|---|---|---|---|---|---|---|---|---|
| A1552 | | 8.9 | 8.9 | 8.9 | 7.5 | 6.7 | 5.7 | 6.7 | 6.7 | 5.3 | 100 | 99.5 | 8.9 | 8.8 | 8.9 | 9.8 |
| SA10G | | | 100 | 100 | 13.9 | 8.4 | 8.2 | 8.4 | 8.4 | 7.4 | 8.9 | 8.9 | 98.9 | 98.9 | 98.9 | 10.4 |
| E7G | | | | 100 | 13.9 | 8.4 | 8.2 | 8.4 | 8.4 | 7.4 | 8.9 | 8.9 | 98.9 | 98.9 | 98.9 | 10.4 |
| SA7G | | | | | 13.9 | 8.4 | 8.2 | 8.4 | 8.4 | 7.4 | 8.9 | 8.9 | 98.9 | 98.9 | 98.9 | 10.4 |
| SL6Y | | | | | | 8.6 | 10.6 | 8.6 | 8.6 | 5.6 | 7.5 | 7.5 | 13.8 | 13.8 | 13.8 | 6.5 |
| SP6G | | | | | | | 9.2 | 100 | 100 | 5.7 | 6.7 | 6.7 | 8.3 | 8.3 | 8.4 | 10.8 |
| L6G | | | | | | | | 9.2 | 9.2 | 5.4 | 5.7 | 5.7 | 8.7 | 8.6 | 8.4 | 8.5 |
| W7G | | | | | | | | | 100 | 5.7 | 6.7 | 6.7 | 8.3 | 8.3 | 8.4 | 10.8 |
| W6G | | | | | | | | | | 5.7 | 6.7 | 6.7 | 8.3 | 8.3 | 8.4 | 10.8 |
| SP7G | | | | | | | | | | | 5.3 | 5.3 | 7.6 | 7.6 | 7.4 | 5.3 |
| W10G | | | | | | | | | | | | 99.5 | 8.9 | 8.8 | 8.9 | 9.8 |
| SA3G | | | | | | | | | | | | | 8.9 | 8.8 | 8.9 | 9.8 |
| SA5Y | | | | | | | | | | | | | | 100 | 100 | 10.8 |
| SL4G | | | | | | | | | | | | | | | 100 | 10.8 |
| SL5Y | | | | | | | | | | | | | | | | 10.9 |
| SO5Y | | | | | | | | | | | | | | | | |

**Large cluster – Immunity**

| | A1552 | SA10G | E7G | SA7G | SL6Y | SP6G | L6G | W7G | W6G | SP7G | W10G | SA3G | SA5Y | SL4G | SL5Y | SO5Y |
|---|---|---|---|---|---|---|---|---|---|---|---|---|---|---|---|---|
| A1552 | | 11.7 | 11.7 | 11.7 | 7.7 | 8.1 | 12.0 | 8.1 | 8.1 | 6.4 | 100 | 100 | 11.7 | 11.7 | 11.7 | 15.8 |
| SA10G | | | 100 | 100 | 10.8 | 6.8 | 8.4 | 6.8 | 6.8 | 10.9 | 11.7 | 11.7 | 100 | 100 | 100 | 10.5 |
| E7G | | | | 100 | 10.8 | 6.8 | 8.4 | 6.8 | 6.8 | 10.9 | 11.7 | 11.7 | 100 | 100 | 100 | 10.5 |
| SA7G | | | | | 10.8 | 6.8 | 8.4 | 6.8 | 6.8 | 10.9 | 11.7 | 11.7 | 100 | 100 | 100 | 10.5 |
| SL6Y | | | | | | 8.5 | 15.8 | 8.5 | 8.5 | 8.8 | 7.7 | 7.7 | 10.8 | 10.8 | 10.8 | 10.1 |
| SP6G | | | | | | | 7.0 | 100 | 100 | 8.4 | 8.1 | 8.1 | 6.8 | 6.8 | 6.8 | 9.7 |
| L6G | | | | | | | | 7.0 | 7.0 | 8.5 | 12.0 | 12.0 | 8.4 | 8.4 | 8.4 | 11.2 |
| W7G | | | | | | | | | 100 | 8.4 | 8.1 | 8.1 | 6.8 | 6.8 | 6.8 | 9.7 |
| W6G | | | | | | | | | | 8.4 | 8.1 | 8.1 | 6.8 | 6.8 | 6.8 | 9.7 |
| SP7G | | | | | | | | | | | 6.4 | 6.4 | 10.9 | 10.9 | 10.9 | 7.5 |
| W10G | | | | | | | | | | | | 100 | 11.7 | 11.7 | 11.7 | 15.8 |
| SA3G | | | | | | | | | | | | | 11.7 | 11.7 | 11.7 | 15.8 |
| SA5Y | | | | | | | | | | | | | | 100 | 100 | 10.5 |
| SL4G | | | | | | | | | | | | | | | 100 | 10.5 |
| SL5Y | | | | | | | | | | | | | | | | 10.5 |
| SO5Y | | | | | | | | | | | | | | | | |

Green: 100% identity (same family and subfamily)
Yellow: 90% - 99.99% identity (same family, with some polymorphism)
Orange: 30% - 90% identity (same family, with considerable polymorphism)
Red: <30% identity (different families)

**Table S5.** Matrices of % identity (percentage of residues that are identical) among the effector and immunity proteins harbored in the T6SS auxiliary cluster 1 from the *V. cholerae* environmental strains and the pandemic A1552 strain.

**Auxiliary cluster 1 – Effector**

| | A1552 | SA10G | E7G | SA7G | SL6Y | SP6G | L6G | W7G | W6G | SP7G | W10G | SA3G | SA5Y | SL4G | SL5Y | SO5Y |
|---|---|---|---|---|---|---|---|---|---|---|---|---|---|---|---|---|
| A1552 | | 15.0 | 15.0 | 15.0 | 15.5 | 15.0 | 15.5 | 15.8 | 15.8 | 15.8 | 14.6 | 15.9 | 15.9 | 15.9 | 14.9 | 15.3 |
| SA10G | | | 100 | 100 | 90.4 | 99.7 | 90.6 | 94.6 | 94.6 | 94.6 | 91.3 | 94.5 | 94.5 | 94.4 | 92.5 | 98.9 |
| E7G | | | | 100 | 90.4 | 99.7 | 90.6 | 94.6 | 94.6 | 94.6 | 91.3 | 94.5 | 94.5 | 94.4 | 92.5 | 98.9 |
| SA7G | | | | | 90.4 | 99.7 | 90.6 | 94.6 | 94.6 | 94.6 | 91.3 | 94.5 | 94.5 | 94.4 | 92.5 | 98.9 |
| SL6Y | | | | | | 90.4 | 99.9 | 92.9 | 92.9 | 92.9 | 87.0 | 92.8 | 92.8 | 92.7 | 94.5 | 91.1 |
| SP6G | | | | | | | 90.6 | 94.4 | 94.4 | 94.3 | 91.3 | 94.2 | 94.2 | 94.1 | 92.3 | 98.9 |
| L6G | | | | | | | | 93.1 | 93.1 | 93.1 | 87.2 | 92.9 | 92.9 | 92.8 | 94.6 | 91.3 |
| W7G | | | | | | | | | 100 | 100 | 90.3 | 99.6 | 99.6 | 99.4 | 91.0 | 94.6 |
| W6G | | | | | | | | | | 100 | 90.3 | 99.6 | 99.6 | 99.4 | 91.0 | 94.6 |
| SP7G | | | | | | | | | | | 90.3 | 99.6 | 99.6 | 99.4 | 91.0 | 94.6 |
| W10G | | | | | | | | | | | | 90.1 | 90.1 | 89.9 | 89.9 | 91.9 |
| SA3G | | | | | | | | | | | | | 100 | 99.9 | 90.8 | 94.5 |
| SA5Y | | | | | | | | | | | | | | 99.9 | 90.83 | 94.5 |
| SL4G | | | | | | | | | | | | | | | 90.7 | 94.4 |
| SL5Y | | | | | | | | | | | | | | | | 92.7 |
| SO5Y | | | | | | | | | | | | | | | | |

**Auxiliary cluster 1 – Immunity**

| | A1552 | SA10G | E7G | SA7G | SL6Y | SP6G | L6G | W7G | W6G | SP7G | W10G | SA3G | SA5Y | SL4G | SL5Y | SO5Y |
|---|---|---|---|---|---|---|---|---|---|---|---|---|---|---|---|---|
| A1552 | | 15.5 | 15.5 | 15.5 | 15.2 | 15.5 | 15.2 | 15.9 | 15.9 | 15.9 | 14.0 | 15.9 | 15.9 | 15.9 | 15.1 | 15.1 |
| SA10G | | | 100 | 100 | 72.1 | 99.6 | 71.7 | 85.7 | 85.7 | 85.7 | 78.3 | 85.7 | 85.7 | 85.7 | 76.2 | 97.5 |
| E7G | | | | 100 | 72.1 | 99.6 | 71.7 | 85.7 | 85.7 | 85.7 | 78.3 | 85.7 | 85.7 | 85.7 | 76.2 | 97.5 |
| SA7G | | | | | 72.1 | 99.6 | 71.7 | 85.7 | 85.7 | 85.7 | 78.3 | 85.7 | 85.7 | 85.7 | 76.2 | 97.5 |
| SL6Y | | | | | | 72.1 | 99.6 | 78.7 | 78.7 | 78.7 | 65.6 | 78.7 | 78.7 | 78.7 | 91.4 | 72.5 |
| SP6G | | | | | | | 71.7 | 85.7 | 85.7 | 85.7 | 77.9 | 85.7 | 85.7 | 85.7 | 75.8 | 97.1 |
| L6G | | | | | | | | 78.3 | 78.3 | 78.3 | 65.2 | 78.3 | 78.3 | 78.3 | 90.9 | 72.1 |
| W7G | | | | | | | | | 100 | 100 | 71.3 | 100 | 100 | 100 | 73.0 | 85.2 |
| W6G | | | | | | | | | | 100 | 71.3 | 100 | 100 | 100 | 73.0 | 85.2 |
| SP7G | | | | | | | | | | | 71.3 | 100 | 100 | 100 | 73.0 | 85.2 |
| W10G | | | | | | | | | | | | 71.3 | 71.3 | 71.3 | 70.2 | 79.2 |
| SA3G | | | | | | | | | | | | | 100 | 100 | 73.0 | 85.2 |
| SA5Y | | | | | | | | | | | | | | 100 | 73.0 | 85.2 |
| SL4G | | | | | | | | | | | | | | | 73.0 | 85.2 |
| SL5Y | | | | | | | | | | | | | | | | 76.3 |
| SO5Y | | | | | | | | | | | | | | | | |

Green: 100% identity (same family and subfamily)
Yellow: 90% - 99.99% identity (same family, with some polymorphism)
Orange: 30% - 90% identity (same family, with considerable polymorphism)
Red: <30% identity (different families)

**Table S6.** Matrices of % identity (percentage of residues that are identical) among the effector and immunity proteins harbored in the T6SS auxiliary cluster 2 from the *V. cholerae* environmental strains and the pandemic A1552 strain.

**Auxiliary cluster 2 – Effector**

| | A1552 | SA10G | E7G | SA7G | SL6Y | SP6G | L6G | W7G | W6G | SP7G | W10G | SA3G | SA5Y | SL4G | SL5Y | SO5Y |
|---|---|---|---|---|---|---|---|---|---|---|---|---|---|---|---|---|
| A1552 | | 14.6 | 14.4 | 14.4 | 98.3 | 94.8 | 14.8 | 94.0 | 94.0 | 14.8 | 96.5 | 94.3 | 93.6 | 93.6 | 93.6 | 12.7 |
| SA10G | | | 95.2 | 95.2 | 14.5 | 15.3 | 97.6 | 15.2 | 15.2 | 97.5 | 14.9 | 15.0 | 15.2 | 15.2 | 15.2 | 20.2 |
| E7G | | | | 100 | 14.3 | 15.0 | 95.1 | 15.0 | 15.0 | 95.0 | 14.6 | 14.7 | 15.1 | 15.1 | 15.1 | 19.8 |
| SA7G | | | | | 14.3 | 15.0 | 95.1 | 15.0 | 15.0 | 95.0 | 14.6 | 14.7 | 15.1 | 15.1 | 15.1 | 19.8 |
| SL6Y | | | | | | 94.1 | 14.7 | 93.7 | 93.5 | 14.7 | 95.7 | 93.8 | 93.7 | 93.7 | 93.7 | 12.8 |
| SP6G | | | | | | | 15.4 | 96.1 | 96.1 | 15.4 | 94.7 | 98.8 | 95.0 | 95.0 | 95.0 | 12.7 |
| L6G | | | | | | | | 15.3 | 15.3 | 99.9 | 15.0 | 15.2 | 15.4 | 15.4 | 15.4 | 19.8 |
| W7G | | | | | | | | | 100 | 15.3 | 93.8 | 94.8 | 99.0 | 99.0 | 99.0 | 13.0 |
| W6G | | | | | | | | | | 15.3 | 93.8 | 94.8 | 99.0 | 99.0 | 99.0 | 13.0 |
| SP7G | | | | | | | | | | | 15.0 | 15.2 | 15.4 | 15.4 | 15.4 | 19.8 |
| W10G | | | | | | | | | | | | 94.0 | 93.4 | 93.4 | 93.4 | 12.8 |
| SA3G | | | | | | | | | | | | | 94.0 | 94.0 | 94.0 | 12.4 |
| SA5Y | | | | | | | | | | | | | | 100 | 100 | 13.0 |
| SL4G | | | | | | | | | | | | | | | 100 | 13.0 |
| SL5Y | | | | | | | | | | | | | | | | 13.0 |
| SO5Y | | | | | | | | | | | | | | | | |

**Auxiliary cluster 2 – Immunity**

| | A1552 | SA10G | E7G | SA7G | SL6Y | SP6G | L6G | W7G | W6G | SP7G | W10G | SA3G | SA5Y | SL4G | SL5Y | SO5Y |
|---|---|---|---|---|---|---|---|---|---|---|---|---|---|---|---|---|
| A1552 | | 13.2 | 12.9 | 12.9 | 99.6 | 72.3 | 12.9 | 68.7 | 68.7 | 12.9 | 83.9 | 71.9 | 68.3 | 68.3 | 68.2 | 12.4 |
| SA10G | | | 73.1 | 73.1 | 13.2 | 13.6 | 92.5 | 12.9 | 12.9 | 92.2 | 12.5 | 13.6 | 12.9 | 12.9 | 12.9 | 9.4 |
| E7G | | | | 100 | 12.9 | 11.2 | 73.9 | 11.9 | 11.9 | 73.9 | 12.2 | 11.5 | 11.9 | 11.9 | 11.9 | 8.1 |
| SA7G | | | | | 12.9 | 11.2 | 73.9 | 11.9 | 11.9 | 73.9 | 12.2 | 11.5 | 11.9 | 11.9 | 11.9 | 8.1 |
| SL6Y | | | | | | 71.9 | 12.9 | 68.3 | 68.3 | 12.9 | 83.5 | 71.5 | 67.9 | 67.9 | 67.9 | 12.4 |
| SP6G | | | | | | | 12.5 | 80.1 | 80.1 | 12.5 | 74.0 | 99.6 | 80.5 | 80.5 | 80.5 | 12.1 |
| L6G | | | | | | | | 12.5 | 12.5 | 99.7 | 12.2 | 12.5 | 12.5 | 12.5 | 12.5 | 9.9 |
| W7G | | | | | | | | | 100 | 12.5 | 67.5 | 79.7 | 99.2 | 99.2 | 99.2 | 11.0 |
| W6G | | | | | | | | | | 12.5 | 67.5 | 79.7 | 99.2 | 99.2 | 99.2 | 11.0 |
| SP7G | | | | | | | | | | | 12.2 | 12.5 | 12.5 | 12.5 | 12.5 | 9.9 |
| W10G | | | | | | | | | | | | 73.6 | 67.5 | 67.5 | 67.5 | 11.2 |
| SA3G | | | | | | | | | | | | | 80.1 | 80.1 | 80.1 | 12.1 |
| SA5Y | | | | | | | | | | | | | | 100 | 100 | 11.0 |
| SL4G | | | | | | | | | | | | | | | 100 | 11.0 |
| SL5Y | | | | | | | | | | | | | | | | 11.0 |
| SO5Y | | | | | | | | | | | | | | | | |

Green: 100% identity (same family and subfamily)
Yellow: 90% - 99.99% identity (same family, with some polymorphism)
Orange: 30% - 90% identity (same family, with considerable polymorphism)
Red: <30% identity (different families)
